# A connectome manipulation framework for the systematic and reproducible study of structure–function relationships through simulations

**DOI:** 10.1101/2024.05.24.593860

**Authors:** Christoph Pokorny, Omar Awile, James B. Isbister, Kerem Kurban, Matthias Wolf, Michael W. Reimann

## Abstract

Synaptic connectivity at the neuronal level is characterized by highly non-random features. Hypotheses about their role can be developed by correlating structural metrics to functional features. But to prove causation, manipulations of connectivity would have to be studied. However, the fine-grained scale at which non-random trends are expressed makes this approach challenging to pursue experimentally. Simulations of neuronal networks provide an alternative route to study arbitrarily complex manipulations in morphologically and biophysically detailed models. Here, we present Connectome-Manipulator, a Python framework for rapid connectome manipulations of large-scale network models in SONATA format. In addition to creating or manipulating the connectome of a model, it provides tools to fit parameters of stochastic connectivity models against existing connectomes. This enables rapid replacement of any existing connectome with equivalent connectomes at different levels of complexity, or transplantation of connectivity features from one connectome to another, for systematic study. We employed the framework in a detailed model of rat somatosensory cortex in two exemplary use cases: transplanting interneuron connectivity trends from electron microscopy data and creating simplified connectomes of excitatory connectivity. We ran a series of network simulations and found diverse shifts in the activity of individual neuron populations causally linked to these manipulations.

## INTRODUCTION

The **structure** of synaptic connectivity decidedly shapes neuronal activity. It can even be said to implement the specific **functions** of different microcircuits. For example, attractor states have been shown to emerge in models with clusters of neurons that are more strongly interconnected than the rest of the population (Deco & Hugues, 2012; Lagzi & Rotter, 2015; Litwin-Kumar & Doiron, 2012). Studying the link between structure and function becomes harder for more complex trends of connectivity, such as overexpression of triad motifs or targeting specificity (Perin, Berger, & Markram, 2011; Pi et al., 2013; Schneider-Mizell et al., 2023; Song, SjÖstrÖm, Reigl, Nelson, & Chklovskii, 2005). Yet, increased complexity that is not captured by comparatively simple connectivity model has been demonstrated to be relevant. For example, clustered inhibition allows competition between attractors without firing rate saturation (Rost, Deger, & Nawrot, 2018), and Renart, Moreno-Bote, Wang, and Parga (2007) speculated that beyond the topology of the wiring diagram, biological details such as dynamic synapses, synaptic failure, dendritic integration, and synaptic clustering may be crucial.

The advent of electron-microscopic (EM) tissue reconstructions, such as the MICrONS dataset (MICrONS Consortium et al., 2021), has been a great boon for researchers studying such important questions, as they provide a complete snapshot of neuronal connectivity instead of sparsely sampled connections. The research to find and describe the mechanisms of connectivity in these datasets that implement the local circuit’s function is still ongoing. However, large-scale EM **connectomes** contain millions of synapses between thousands of neurons, allowing the discovery of any number of non-random trends. But which ones are functionally relevant, and which ones are mere epiphenomena? Co-registered neuron recordings enable their correlation with function, but to demonstrate causation, a change of function must be the result of a manipulation that affects the structural trend observed. Currently, the only viable way to conduct such an experiment is ***in silico***. Recently, modeling techniques have been developed that include sufficient biological detail to reproduce non-random connectivity trends observed in biology (Billeh et al., 2020; Isbister et al., 2024; Markram et al., 2015). An even more powerful approach could be based on ***transplanting*** the connectivity observed in EM into an *in silico* model. Both cases can then be followed by *connectome manipulations* that add or remove connectivity trends, together with observations of their functional impact.

While promising, the outlined research faces confounding factors and particular difficulties: activity *in silico* is affected by simplifications and assumptions that are inherent to the process of modeling. As such, any *in silico* model should be carefully validated against experimental data, and the effect of connectome manipulations should be probed in different baseline models to assess robustness of the results. Other challenges arise from the fact that detailed, bottom-up models include multi-synaptic connectivity (Reimann, King, Muller, Ramaswamy, & Markram, 2015), i.e., multiple and individually parameterized synapses forming a connection from one neuron to another. This usually leads to large data sizes and complicated file structures that are necessary to store connectivity at such a level of detail. Hence, manipulations must not only be efficient enough to deal with large amounts of data but also preserve biological distributions of parameter values and respect known biological principles, e.g., Dale’s law (Strata & Harvey, 1999). Additionally, non-random trends of interest exist in the higher-order structure of connectivity (Perin et al., 2011; Song et al., 2005) or at the level of subcellular targeting of **pathways** (Pi et al., 2013; Schneider-Mizell et al., 2023). Therefore, specifically manipulating them while keeping the connectivity otherwise unchanged can be conceptually or mathematically challenging.

Here, we present Connectome-Manipulator, a programmatic Python framework that enables the creation, transplantation, and manipulation of connectomes. Based on the Scalable Open Network Architecture TemplAte (SONATA) standard (Dai et al., 2020), the framework allows manipulations to be applied to any network model described in that standard, while their functional impact can be readily investigated through network simulations using any simulator supporting SONATA. Notably, SONATA is a format for multiscale neural network models as well as their simulation outputs and was designed for memory and computational efficiency and cross-platform compatibility. It is based on a non-stochastic representation of a network that explicitly includes multi-synaptic connections by storing each individual synapse and its parameters together with its pre- and post-synaptic neuron. This allows the representation of network models at any level of topological complexity as well as connectome reconstructions from EM. On the one hand, we offer as part of the framework algorithms to systematically simplify the connectivity of a given SONATA model. While simplifying the structure of the wiring diagram, other anatomical and physiological parameters are carefully preserved. This allows the user to start with any proposed complex connectome and then study the impact of simplification. On the other hand, we also provide functionality to add new complexity to a connectome, as defined by an arbitrarily complex **adjacency** or **synaptome matrix**.

While an adjacency matrix represents the connectivity between pairs of neurons, a synaptome matrix can be used to even define the exact number of synapses that are part of a given connection, thereby offering great flexibility in studying the effect of specific manipulations while preserving other aspects of connectivity.

In addition, while not the main focus, the framework also enables basic manipulations, such as adjustments of physiological synapse parameters, or specific removal of synapses or connections as in lesion experiments. While both SONATA and our framework were developed with morphologically and physiologically detailed models in mind, they also support point neuron models. Thus our framework can be used for example to transplant a wiring diagram from a detailed to a point neuron model, in order to study manipulations on that level.

We demonstrate the applicability of our framework by manipulating the connectome of a detailed model of the rat somatosensory cortex (Isbister et al., 2024; Reimann et al., 2024) in two particular ways. First, we increased inhibitory targeting specificity of vasoactive intestinal peptide-expressing (VIP+) interneurons, thereby transplanting connectivity trends present in the MICrONS dataset (Schneider-Mizell et al., 2023). We found that despite the fact that the VIP+ interneurons are predominantly targeting other inhibitory neurons in the manipulated connectome, their activation can still lower the firing rate of excitatory populations, but to a lesser extent than in the original connectome. Second, we studied how progressively simplified (Gal, Perin, Markram, London, & Segev, 2020), but otherwise equivalent connectivity among excitatory neurons affects the dynamics of spiking. We found that layer 4 excitatory neurons were consistently shifted towards a more internally driven spontaneous activity regime, while layer 6 inhibitory neurons were shifted towards a more externally driven regime. For layer 6 excitatory neurons we found diverse effects depending on the degree of simplification. Taken together, these results demonstrate that our framework allows emergent network activity to be causally linked to specific features of connectivity in a systematic and reproducible way.

## RESULTS

### Connectome Manipulation Framework

The Connectome-Manipulator software presented in this manuscript is a universal framework for creating and manipulating connectivity in large-scale models of neural networks in SONATA format (Dai et al., 2020). Its source code is openly available on GitHub under https://github.com/BlueBrain/connectome-manipulator. Manipulations can be applied to entire models, specific sub-networks, or even single neurons, ranging from insertion or removal of specific motifs to complete **rewiring** based on **stochastic connectivity models** at various levels of complexity. Important scientific use cases include **wiring** a connectome from scratch based on given connectivity rules, rewiring an existing connectome while preserving certain aspects of connectivity, and transplanting specific connectivity characteristics from one connectome to another (Figure 1A).

**Figure 1.**
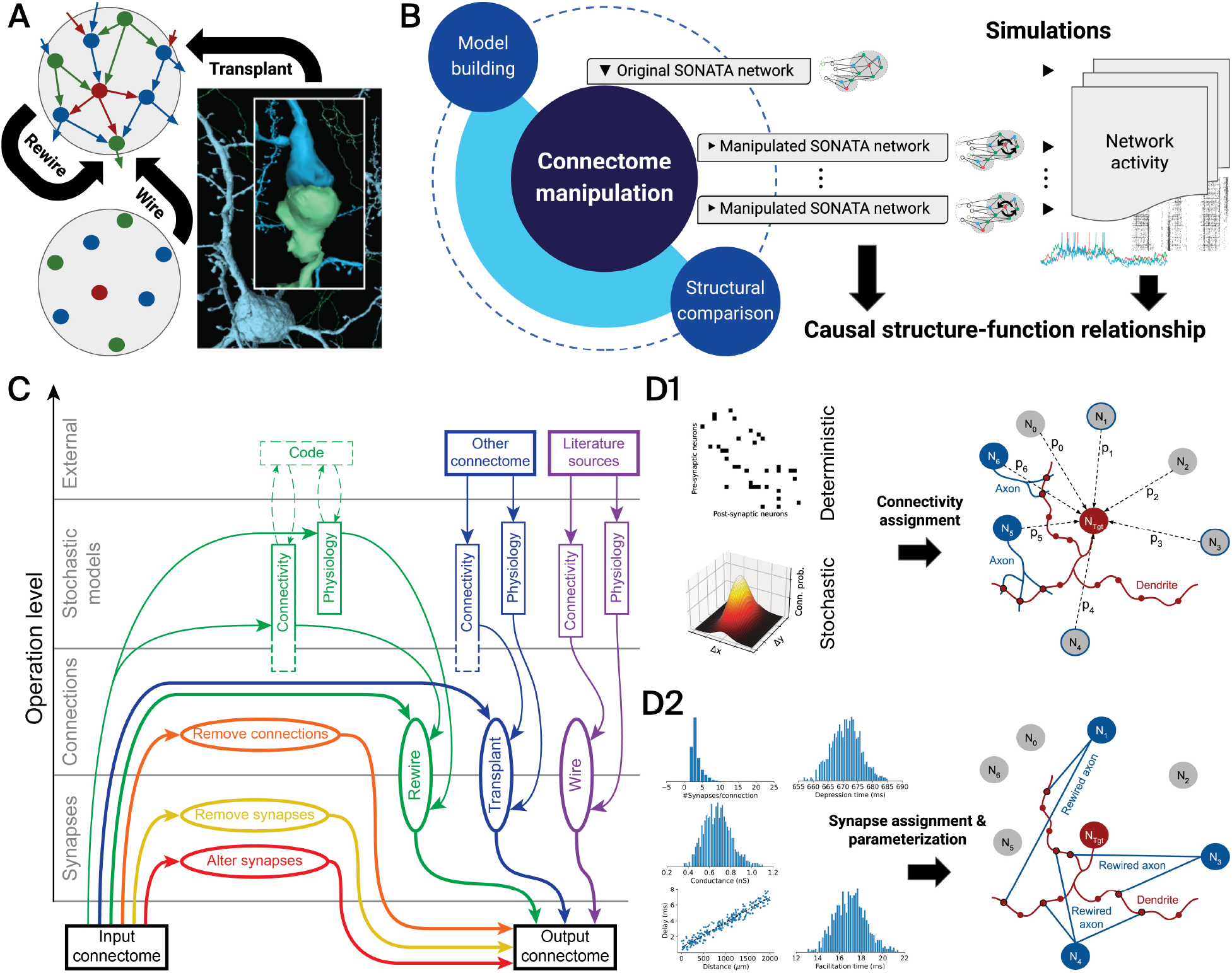
Connectome manipulation framework. (A) Scientific use cases, such as wiring (generating) or rewiring (changing) a connectome based on connectivity rules, or transplanting specific connectivity characteristics from other connectomes (e.g., from EM reconstructions from MICrONS Consortium et al. (2021) as shown here). (B) Main components of the framework, including functionality for connectome manipulations, model building (i.e., fitting stochastic model representations against existing connectomes), and structural comparison of manipulated connectomes (cf. Figure 2). A typical experimental workflow involves a series of connectome manipulations (e.g., with different levels of severity) of a given original SONATA network model, resulting in a set of manipulated SONATA network models. Computer simulations of the original and manipulated network models can then be run in order to causally link the emergent network activity to certain structural features of connectivity. (C) Operation levels and how they interact in six typical connectome manipulation scenarios, as indicated by color. Alteration and removal of individual synapses operate at the level of synapses; removal of entire connections at the level of connections; rewiring, transplantation, and wiring at both of these levels. In some of these scenarios, stochastic model descriptions for connectivity and/or physiology are required, which may be derived from the input connectome or from external sources. (D) Algorithmic steps for establishing new connections: first, connectivity assignment, supporting deterministic and stochastic descriptions of connectivity and turning them into a connectome at the level of connections (D1); second, synapse assignment and physiological parameterization, allowing pathway-specific parameter distributions, for expanding the connectome into a description at the level of synapses (D2).

The main components of the framework include functionality for connectome manipulation, model building, i.e., fitting a stochastic connectivity model against an existing connectome, and structural comparison of manipu- lated connectomes (Figure 1B). The connectome manipulation functionality applies one or a sequence of manip- ulations to a given connectome of a network model in SONATA format. In this format, nodes (i.e., neurons) and edges (i.e., connections, formed by synapses) are represented as table-based data structures in Hierarchical Data Format version 5 (HDF5) format. Specifically, the node and edge tables contain the respective properties of each individual neuron and synapse. While SONATA has predefined properties and naming conventions, the format is loosely defined in the sense that it allows the definition of user-defined node and edge properties. Accordingly, our framework can deal with various properties as defined by the user in a flexible way. Manipulations can be targeted to the entire connectome or to selected pathways, i.e., connections between specific pre- and post-synaptic neurons, based on criteria such as their morphological type (m-type) or electrophysiological type. The output of such manipulation(s) is again a network model in SONATA format consisting of the same set of neurons as the original network, together with the manipulated connectome. A typical experimental workflow starts with the creation of a set of manipulated connectomes (e.g., with different levels of severity) from a given baseline connectome. The resulting connectomes can be readily simulated using any simulator supporting SONATA, allowing the systematic and reproducible characterization of causal effects of structural manipulations on the network activity and function.

Conceptually, a biophysically detailed connectome can be described on different levels of detail (Figure 1C): The *synapse* level preserves the full level of detail, describing individual synapses and their anatomical and physiological parameters. The *connection* level simplifies this to an adjacency matrix, i.e., representing whether or not a connection from one neuron to another exists. The *stochastic model* level simplifies this further to a stochastic description, for example a notion of distance-dependent connection probability. Note that within this level, various types of stochastic model descriptions with different amounts of detail exist (see Figure 2B and Table S1). At this level, physiological parameters are also stochastically described by probability distributions. This leads to six classes of connectome manipulations that differ in the levels of descriptions they employ, and where manipulations are applied to (Figure 1C):

**Figure 2.**
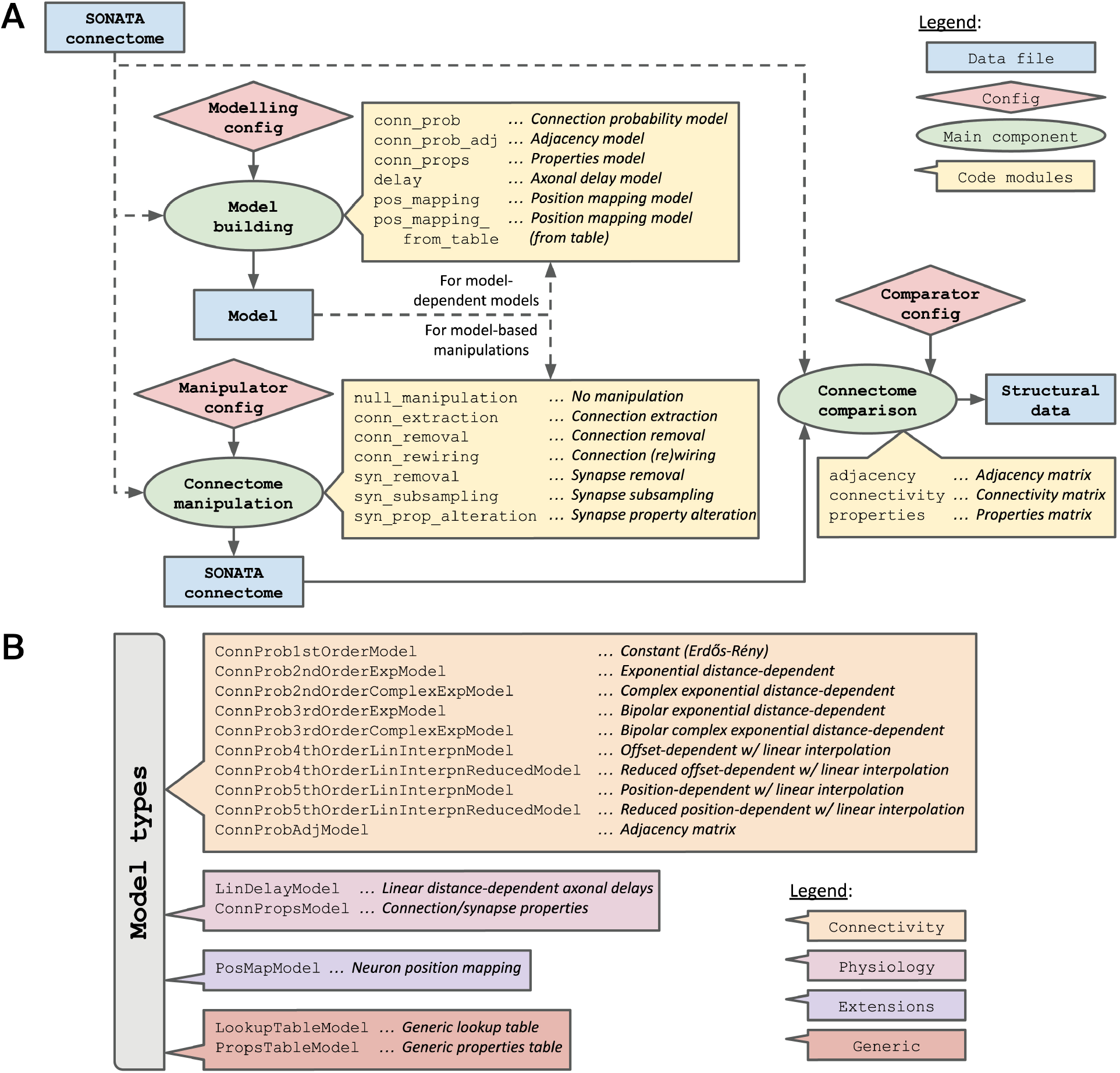
Software reference for the connectome manipulation framework. (A) Main software components for model building, connectome manipulation, and structural comparison and how they are intercon- nected. Additional elements, such as configurations, input/output data files, and their respective code modules are illustrated as indicated by the legend. Dashed lines denote optional interconnections dependent on the actual use case (cf. Figure 1C). More details about the individual code modules can be found in the supplementary reference Tables S2, S3, and S6, as well as in the software documentation (including a configuration file reference). (B) Available model types implemented in the connectome manipulation framework as Python classes under `/model building/model types.py`. The models are divided into different categories as indicated by the legend. More details about the individual model types can be found in the supplementary reference Table S1.

**Alter synapses (red):** Keeps the description on the level of synapses and changes physiological parameters, such as conductance.

**Remove synapses (yellow):** Keeps the description on the level of synapses but removes some of them according to specified selection criteria of pre- and post-synaptic neurons.

**Remove connections (orange):** Works similarly but on the level of connections, i.e., removing entire connections according to specified selection criteria of pre- and post-synaptic neurons.

**Rewire (green):** Rewires an existing connectome, that is, creating new synapses based on a description on the level of connections or stochastic models. Existing synapses may be kept or removed. Additional adjustments can optionally be performed manually at this stage or be implemented in external programming code (dashed lines). This can be done independently for the structure and physiology of connections.

**Transplant (blue):** Same as rewiring, but the source of the connectivity and/or physiological description is another connectome, allowing the transfer of structural and/or physiological characteristics of single connections, pathways, or entire connectomes from one to another connectome.

**Wire (purple):** Same as rewiring, but the input connectome is empty and the connectivity and physiological descriptions are derived from experimental data or literature.

At the end of a connectome manipulation an output at the level of synapses is generated. Simplifying a connectome to the level of a stochastic model will usually drastically alter many aspects of connectivity, such as dendritic locations of synapses from different sources, or in-degree distributions. Various options exist in order to carefully preserve such aspects during this process (see Tables S4 and S5).

For generating new synaptic connectivity when rewiring, transplanting, or wiring connectivity by use of the `conn rewiring` operation (see Section: *Software Architecture of the Connectome Manipulation Framework*), an instance at the level of connections is built first in the “connectivity assignment” step (Figure 1D1). During this step, the pre-synaptic neurons that are to be connected to each post-synaptic neuron are assigned. This is done by determining the connection probability *p*_*i*_ of all potential pre-synaptic source neurons *N*_*i*_ to be connected with a given post-synaptic target neuron *N*_*Tgt*_. The connection probabilities are obtained from a deterministic or stochastic connection probability model (Table S1). Then, new source neurons are randomly sampled from all *N*_*i*_ according to *p*_*i*_ either as independent Bernoulli trials (default), or by optionally preserving the in-degree (`keep indegree` option; Tables S4 and S5) by sampling exactly the same number of incoming connections as in the original connectome independently for each post-synaptic neuron. If the sampled connections already existed in the original connectome, their synapse assignment and parameterization can optionally be kept unchanged; otherwise, they will be replaced by reusing other existing or generating and parameterizing new connections (`keep conns` and `reuse conns` options; Tables S4 and S5). In case the provided connection probability model returns only probability values zero and one, the resulting connectivity assignment will be deterministic.

The level of connections is then further expanded into a description at the level of synapses in the “synapse assignment and parameterization” step (Figure 1D2). During this step, synapses on the post-synaptic dendrite are placed, parameterized, and (randomly) assigned to form all incoming connections as determined in the first step, unless existing connections are kept or reused (Table S5). New synapses can be placed by either duplicating existing synapse positions, randomly generating new positions on the dendrite, or loading positions externally (`reuse`, `random`, and `external` options; Tables S4 and S5). The number of synapses to assign to a new connection, as well as their physiological parameter values, can be sampled from existing connections or are given by a stochastic `ConnPropsModel` (Table S1) defining their (pathway-specific) parameter distributions (`sample` and `randomize` options; Tables S4 and S5). Alternatively, the number of synapses per connection can also be provided deterministically through a synaptome matrix. By use of a `ConnPropsModel`, physiological parameter values are by default drawn independently. An option exists to define correlations between selected synapse parameters by means of pathway-specific covariance matrices (see Table S1 for details).

### Software Architecture of the Connectome Manipulation Framework

The connectome manipulation framework has three main software components for connectome manipulation, model building, and connectome comparison respectively (Figure 2A) which can be invoked through a command line interface. Each tool can be configured by means of a configuration file in JavaScript Object Notation (JSON) format, the structure of which is documented in the software documentation (see Section: *Software and Data Availability*). All manipulation, model building, and comparison functions are implemented in separate Python code modules based on reusable primitives, which makes it easy to add new functionality to the framework (see *Extensions* in Table 1). A variety of connectome manipulation operations as well as functions for model building and structural comparison already exist for different use cases (Figure 2A, yellow boxes). Specifically, various operations belonging to the six classes of manipulations mentioned before (cf. Figure 1C) are available, such as `syn prop alteration` for altering synapses, `syn removal` and `syn subsampling` for remov- ing synapses, `conn extraction` and `conn removal` for removing connections, and `conn rewiring` for rewiring, transplanting and wiring connectivity.

**Table 1.** Useful resources for the connectome manipulation software. This table summarizes the available resources that will be helpful for a user to get started and learn how to use the connectome manipu- lation software.

| Topic | Description | Resources |
| --- | --- | --- |
| Installation | Install instructions for the Python package | README file (GitHub) |
| Usage | Launch commands of the command line tools | README file (GitHub) |
| Error handling | Notes on how to avoid and deal with common errors | README file (GitHub) |
| Code modules | Overview and description of the available code modules | Figure 2; Tables S1-S6 |
| API* reference | Methods reference and description of their parameters | Documentation (Read the Docs) |
| Config files | Reference for respective configuration file structures | Documentation (Read the Docs) |
| Basic examples | Simple examples of some of the features of the framework | Examples folder (GitHub) |
| Advanced examples | Manipulation experiments described in this manuscript | SSCx repository (GitHub) |
| Contributions | Guidelines on how to suggest new features, report bugs, etc. | Contribution guide (GitHub) |
| Extensions | Outline of how to implement new features in the framework | Contribution guide (GitHub) |

\*API ... Application programming interface

Connectome comparison functions allow a user to compare connectivity-related properties of two connectomes. Specifically, they allow the comparison of the baseline with a manipulated connectome with regard to differences in their connectivity structure and their distributions of synaptic properties, together with visualizations on the single-neuron level or grouped by populations of neurons.

A *model* in the context of this framework is a simplified (stochastic) representation of certain aspects of connectivity required by some of the manipulation functions, e.g., connection probability at a given distance (Figure 2B). Models can be stored as JSON files and are thus human-readable and editable; some models containing large amounts of data have them stored in an additional HDF5 file for easy machine processing. The process of *model building* refers to fitting models against existing connectomes and producing a model file. The idea of model fitting is to capture certain aspects of an existing connectome, for example physiological parameter distributions, such that during rewiring, i.e., when generating new connections, their parameterization provided by such a model will be consistent with the original connectome (unless the parameterization itself is subject to manipulation). Since the synapse physiology in biology is known to vary with pre- and post-synaptic cell types (see list of references in Markram et al. (2015)), the stochastic model of type `ConnPropsModel` is able to capture individual physiological parameter distributions for all pairs of pre- and post-synaptic m-types. Other than that, the model fitting tools do not impose any strong biological constraints on the fitted models; it is left to the user to make choices (e.g., about the underlying types of parameter distributions) and determine their biological validity depending on the actual use case. When fitting a model, validation plots are automatically generated which should aid the user in determining the goodness of fit as well as validating the model. Optionally, stochastic models can be fitted with cross-validation in order to prevent overfitting, i.e., fitting a model too closely to a specific dataset instead of capturing the underlying biological principle. (see Methods: *Model Fitting with Cross-Validation*). Some generic model formats and extensions to existing models also exist, allowing for a more fine-grained control (e.g., fitting connectivity models with position mapping).

A detailed description of all the above-mentioned software components can be found in the supplementary Tables S1-S6. Finally, Table 1 provides an overview of useful resources and where to find them, such as install, usage, and error handling instructions, but also application programming interface (API) references, and examples. These resources will help a user to get started with using the connectome manipulation software, extending it if needed, and potentially contributing to our software project.

### Operation Principle and Performance of Connectome Manipulations

Connectome manipulations follow a block-based operation principle and can be run either serially or in parallel on multiple computing cores or nodes (Figure 3A). This works by splitting the input SONATA connectome post- synaptically into disjoint edge tables each of which contains all synapses targeting a block of post-synaptic neurons, the size of which can be configured. Each edge table is then manipulated independently, and the output is written to a separate .parquet file (i.e., in Apache Parquet format). After all manipulations are completed, the individual .parquet files can be kept for further processing, and/or can be merged to a single output SONATA connectome using the parquet-converters utilities (external dependency). Note that the same operation principle also applies for wiring an empty connectome from scratch, in which case all input edge tables will be initialized empty and each manipulation will add new edges to them. Importantly, during connectome manipulations, the node tables of the network model are always kept unchanged. Also, in SONATA, nodes of a network model can be arranged into multiple neuron populations (e.g., representing different brain regions). Similarly, the connectome can consist of multiple edge populations each of which has a specific source and target neuron population. Accordingly, our framework supports network models with multiple node and/or edge populations, but only one selected edge population can be used in a single run. Therefore, where applicable, the user has to specify the respective node and/or edge population names to be used for model fitting, manipulation, or structural comparison runs.

**Figure 3.**
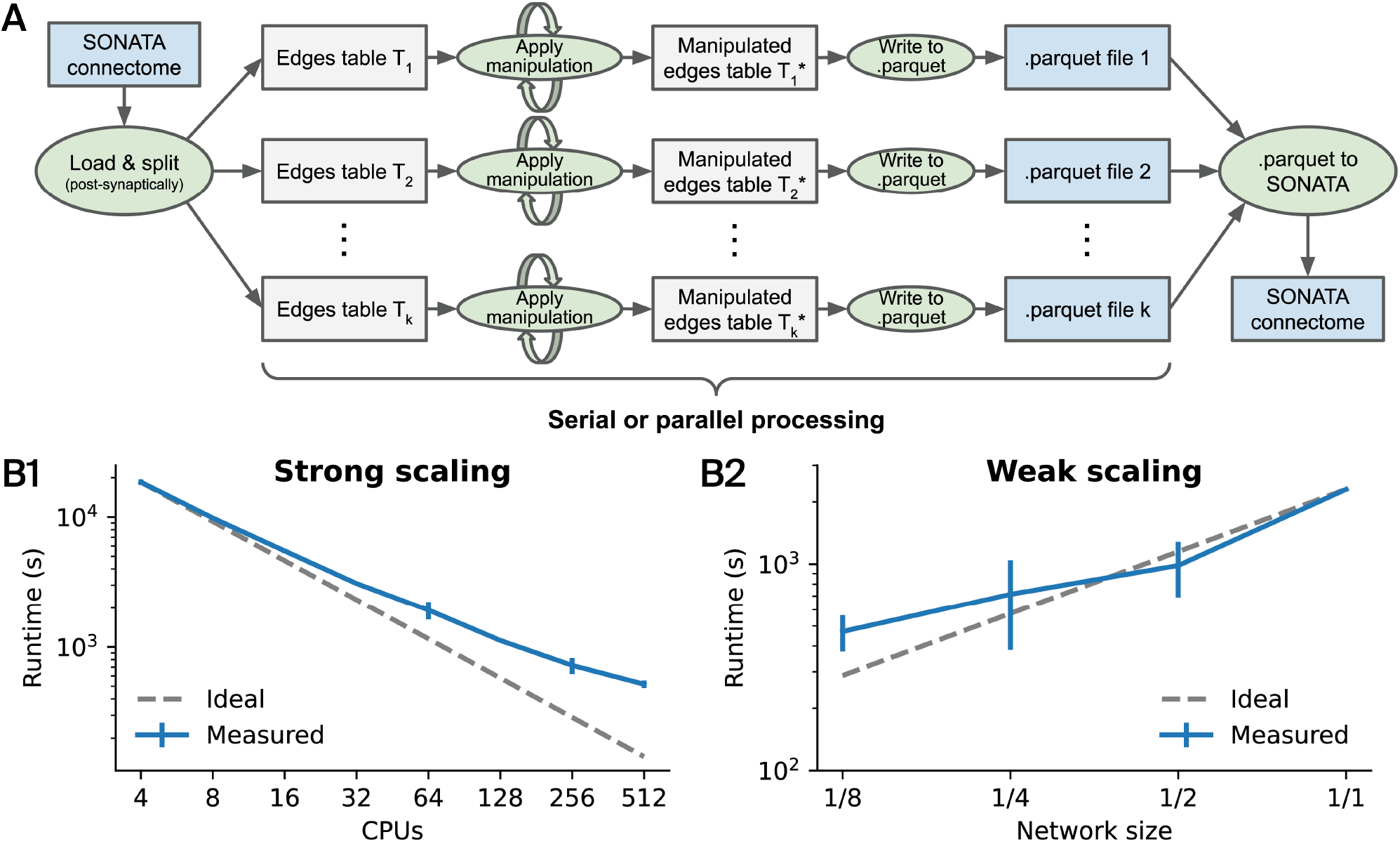
Operation principle and performance of connectome manipulations. (A) Internally, a connectome manipulation works by splitting the input SONATA connectome by post-synaptic neurons into disjoint edge tables. Each table is manipulated independently, and the output is written to a separate .parquet file. This enables operations to be run in series or in parallel. After all manipulations are completed, the individual .parquet files are merged to a single output SONATA con- nectome using parquet-converters (external dependency). (B) Performance evaluation when running rewiring (i.e., `conn rewiring` operation) of the connectivity between excitatory neurons in the central column of a detailed model of the rat somatosensory cortex (as in Figure 4A) based on a 1^st^ order model of connectivity (as in Figure 5A). We ran the same operation with different numbers of processing units (CPUs) in parallel and measured the runtime, resulting in a strong scaling plot (B1). We also ran the same operation with a fixed number of 32 CPUs in parallel but with different network sizes (i.e., random subsets of neurons, as indicated by the fraction), resulting in a weak scaling plot (B2). Error bars denote the standard deviation over two (B1) and four (B2) measurements respectively. Gray dashed lines indicate the ideal scaling behavior without any overhead. Note that both plots are in a double logarithmic scale.

Notably, SONATA as well as our framework also support point neuron populations without detailed morphologies in which case synapses don’t have properties related to afferent synapse locations on the post-synaptic dendrites. Accordingly, when rewiring such a connectome, the morphology-related synapse position options `randomize` and `external` for randomized and externally loaded synapse positions on the dendrites respectively (see Table S4) are not applicable. Also, wiring such a connectome from scratch is currently not supported as it will by default be initialized with afferent location properties. Specifically, for placing new synapses on detailed morphologies, the afferent properties “afferent section id” (section index of the dendritic morphology; each section consists of one or more segments), “afferent section pos” (normalized position offset within a section), “af- ferent section type” (type index: 1 – soma, 3 – basal dendrite, 4 – apical dendrite) as well as “afferent center x”, “… y” and “… z” (x, y, z positions along the axis of the associated segment) will be generated. More details about morphologies can be found in the SONATA developer guide and the MorphIO documentation.

Our connectome manipulation framework has been developed and optimized for large-scale models of neural networks. Specifically, we make great use of bluepysnap and libsonata for accessing SONATA files, of MorphIO and NeuroM for accessing detailed morphologies, as well as of pandas data frames and NumPy arrays together with their integrated operations when generating and processing the huge edge tables that contain the properties of individual synapses. Potential bottlenecks are input/output (I/O), in particular when access to detailed morphologies is required for placing new synapses, as well as working memory. However, the memory bottleneck can be circumvented: By use of *N* data splits, the memory consumption on a single computation node can potentially be reduced by a factor of *∼N* while the runtime should not be affected, except for some overhead.

We theoretically assessed the asymptotic algorithmic complexity of the different manipulation operations that are currently available in the framework (as in Figure 2 and Table S3) based on the “Big O notation” (Bachmann, 1894) using two variables, the number of nodes *n* and the number of edges *e* (Table 2). While the overall I/O overhead is of order *O*(*e*), we found that most manipulation operations have an order *O*(*e* log *e*) which is mainly due to internal synapse selection, grouping, and sorting operations. The complexity of the connectome (re)wiring operation `conn rewiring` can be divided into the complexities of the different algorithmic steps: connectivity assignment *O*(*n*^2^), synapse parameterization *O*(*e*), plus some overhead *O*(*e* log *e*). Overall, under the assumption that *O*(*e*) *≈ O*(*n*^1.4^) in biological neural networks (Azevedo Carvalho, Contassot-Vivier, Buhry, & Martinez, 2020) the whole term can be approximated by *O*(*n*^2^), i.e., the asymptotic complexity is determined by connectivity assignment as it involves assigning connections between all pairs of neurons.

**Table 2.** Algorithmic complexity of the different manipulation operations. We use the “Big O notation” (Bachmann, 1894) to theoretically assess the asymptotic algorithmic complexity based on two variables: *n* (number of nodes) and *e* (number of edges). The connectome (re)wiring operation is divided into algorithmic steps. Details about all manipulation operations can be found in the supplementary reference Table S3.

| Operation | Description | Complexity |
| --- | --- | --- |
| Base framework | Mainly I/O overhead | $O(e)$ |
| null_manipulation | No actual manipulation | $O(1)$ |
| syn_subsampling | Synapse subsampling | $O(e)$ |
| syn_removal | Synapse removal | $O(e \log e)$ |
| syn_prop_alteration | Synapse property alteration | $O(e \log e)$ |
| conn_extraction | Connection extraction | $O(e \log e)$ |
| conn_removal | Connection removal | $O(e \log e)$ |
| conn_rewiring | Connection (re)wiring | $O(n^2)^*$ |
| | ◊ Connectivity assignment | $O(n^2)$ |
| | ◊ Synapse parameterization | $O(e)$ |
| | ◊ Overhead | $O(e \log e)$ |

\* $O(n^2) + O(e) + O(e \log e) \approx O(n^2)$ in biological neural networks, assuming $O(e) \approx O(n^{1.4})$ (Azevedo Carvalho et al., 2020)

Finally, we evaluated the performance of the framework by running a series of benchmark tests. For this purpose, we ran rewiring of the connectivity between excitatory neurons in the central column of a detailed model of the rat somatosensory cortex (as in Figure 4A) based on a stochastic 1^st^ order model of connectivity with a constant connection probability (as in Figure 5A) in two particular ways. First, we assessed strong scaling behaviour, which is a measure of how the computation time scales with different numbers of processing units (CPUs) for a fixed problem size. We therefore ran the same rewiring operation with an arbitrary (large) number of 12,345 data splits using 4 up to 512 CPUs in parallel and measured the required runtime (Figure 3B1). We found good strong scaling behavior for lower CPU counts but an increasing effect of the non-computational overhead for higher CPU counts. This means that most of the runtime is spent on setting up the actual computation in this case. Second, we assessed weak scaling behaviour as how the computation time scales with different problem sizes but a fixed number of CPUs. We therefore ran the same rewiring operation with an again arbitrary number of 1,234 data splits using 32 CPUs in parallel on different network sizes which we obtained from pre-defined node sets containing random subsamples of a given fraction of the 30,190 neurons in the central column of the full network model (Figure 3B2). We found good weak scaling behavior for large network sizes but suboptimal performance for smaller problem sizes as the available resources are less efficiently used in this case. The total memory usage was found to be between 134 GB (full size) and 128 GB (1/8 size), which corresponds to only *∼*4 GB per CPU.

**Figure 4.**
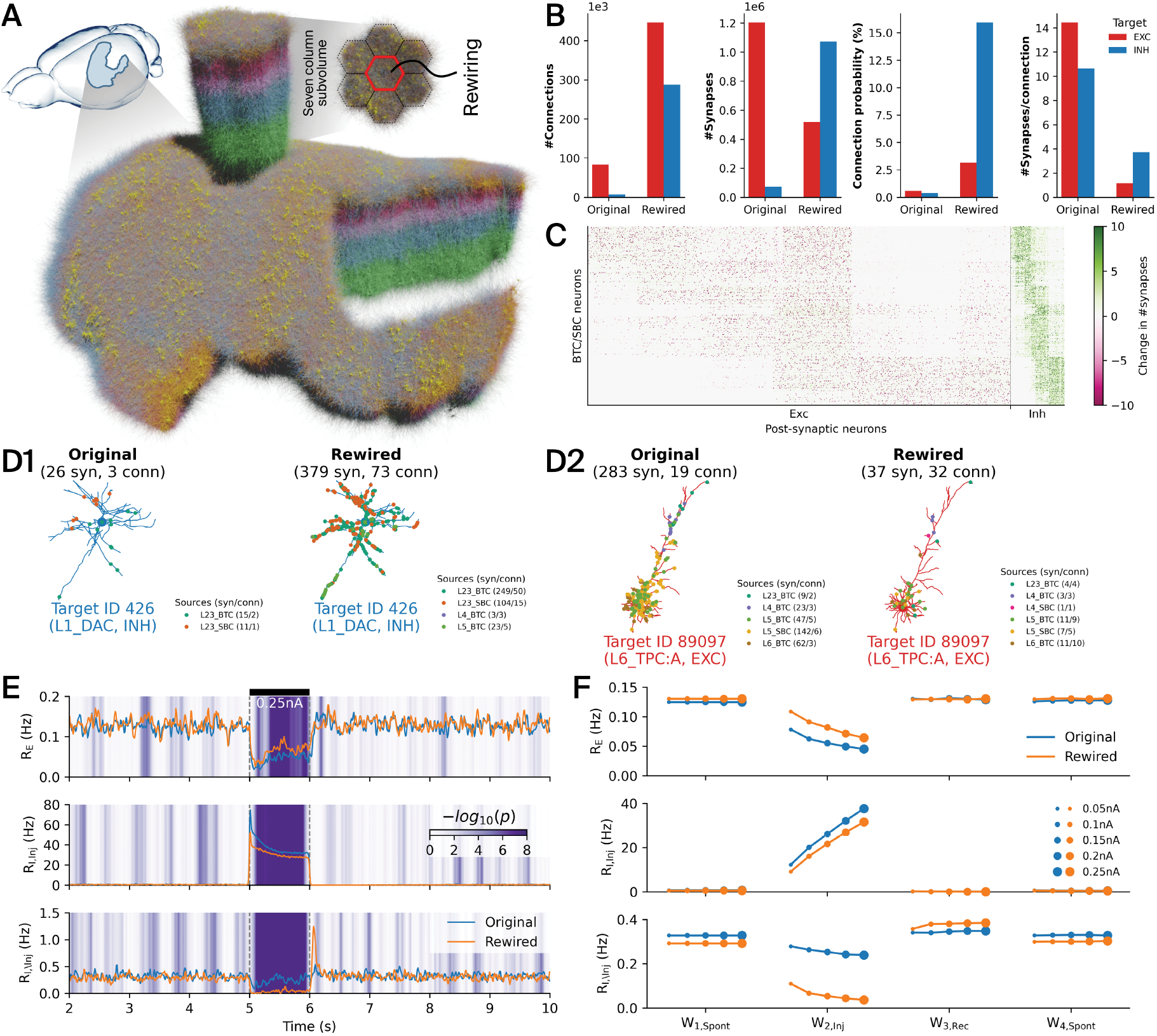
Rewiring of VIP+ interneuron connectivity in a detailed model of rat somatosensory cortex. (A) Large-scale, anatomically (Reimann et al., 2024) and physiologically (Isbister et al., 2024) detailed model of the rat non-barrel somatosensory cortex. The central cortical column (red hexagon) of the seven column subvolume was subject to rewiring. See details in Methods: *Detailed Network Model of the Rat Somatosensory Cortex*. (B) Total number of connections, synapses, mean connection probability and mean number of synapses per connections in the original vs. the rewired connectome between VIP+ source neurons, i.e., bitufted cells (BTC) and small basket cells (SBC), and excitatory (red) and inhibitory (blue) target neurons. (C) Change in numbers of synapses per connection after rewiring, showing all pairs of 530 source (BTC and SBC types; y-axis) and 30,190 target neurons (26,787 excitatory, 3,403 inhibitory; x axis). Only 10 % of the actual density is plotted. The color scale spans the *±* 90^th^ percentile of the values. These results were obtained by running a structural comparison using the `adjacency` code module (see Table S6). (D) Synapses from different BTC/SBC source neurons (as indicated by the legend) targeting an exemplary inhibitory (D1) and excitatory (D2) neuron in the original (left) vs. the rewired (right) connectome. Small numbers in parentheses denote numbers of synapses and connections respectively. More examples can be found in Figure S3 and S4. (E) Instantaneous firing rates during a current injection experiment (estimated in 10 ms bins; smoothed with Gaussian kernel with standard deviation 1.0) of excitatory (*R*_*E*_) and inhibitory (*R*_*I,Inj*_ : BTC/SBC types, injected with current; *RI, \ Inj* : other inhibitory types, not injected with current) populations. A constant 0.25 nA current was injected into BTC/SBC types from time point 5 s to 6 s, as indicated by the black bar. The background shading denotes significant differences between original and rewired activity, computed as the negative decimal logarithm of the p-value of a Wilcoxon rank-sum test applied on samples from a 200 ms sliding window. (F) Average firing rates of excitatory and inhibitory populations for all injection currents (0.05 to 0.25 nA) and different time windows before (*W*_1,*Spont*_), during (*W*_2,*Inj*_), when recovering from (*W*_3,*Rec*_), and after (*W*_4,*Spont*_) current injection (see Methods).

**Figure 5.**
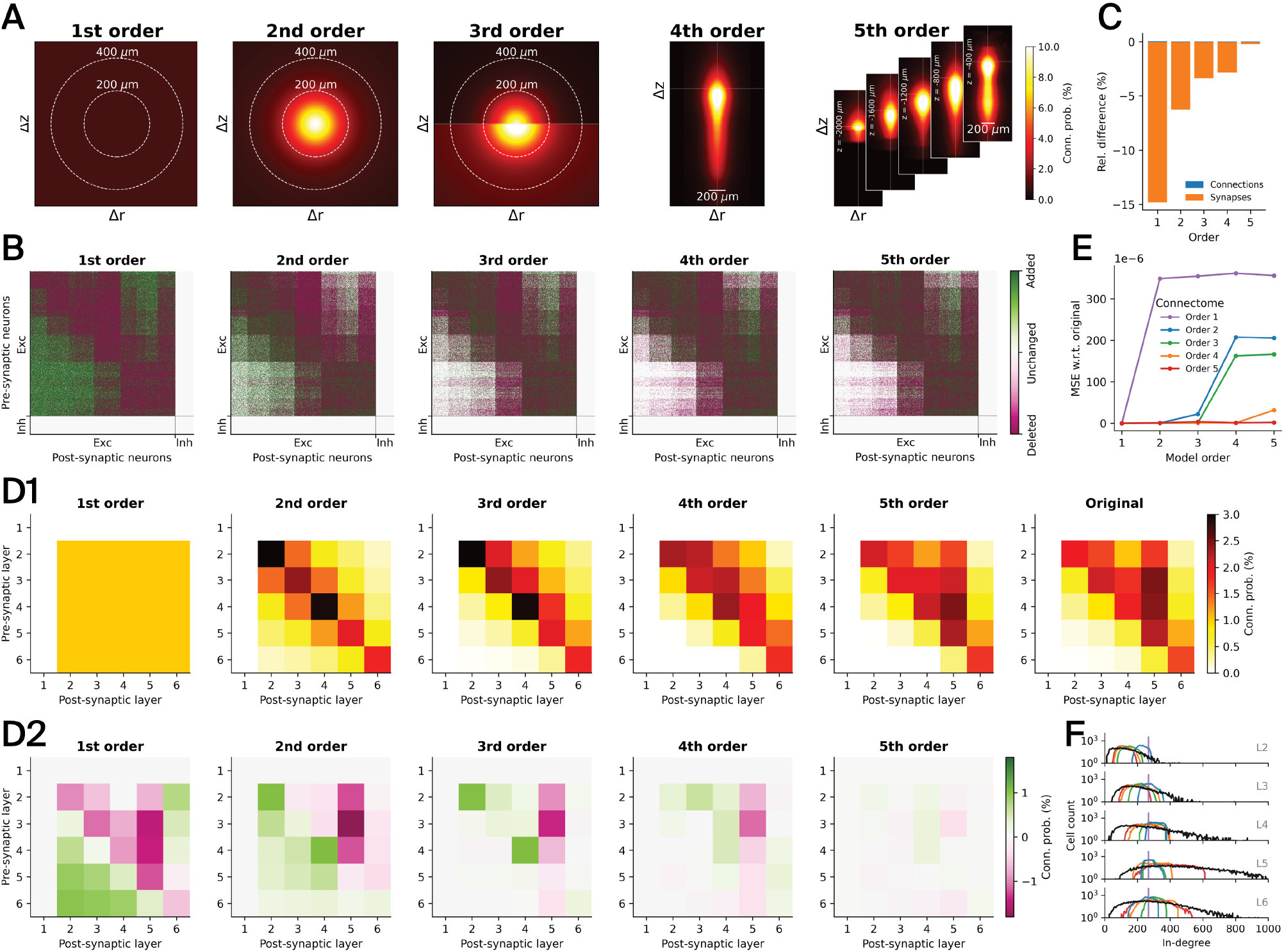
Simplified connectivity of a detailed model of rat somatosensory cortex. (A) All connections between excitatory neurons in the central cortical column of the model were rewired based on five simplified stochastic models of connectivity, whose parameters were fitted against the actual connectivity data from the detailed cortical model: 1^st^ order — constant, 2^nd^ order — distance-dependent, 3^rd^ order — bipolar distance-dependent, 4^th^ order — offset-dependent, and 5^th^ order — position-dependent (Δ*r*… radial offset, Δ*z*… axial offset, *z*… axial position). (B) Resulting adjacency matrices after rewiring, indicating deleted, added, and unchanged connections relative to the baseline connectome. Neurons are order by cell type (excitatory, inhibitory) and layer. Only 10 % of the actual density is plotted. These results were obtained by running structural compar- isons using the `adjacency` code module (see Table S6). (C) Relative differences of the numbers of synapses and connections between excitatory neurons with respect to the baseline connectome (see Table 4 for exact numbers). (D) Average connection probabilities be- tween excitatory neurons in different layers (D1), and differences to baseline connectome (D2). Note that layer 1 does not contain any excitatory neurons. These results were obtained by running structural comparisons using the `connectivity` code module (see Ta- ble S6). (E) Mean squared error of the connection probabilities obtained from the given stochastic models (x-axis) with parameters fitted against the simplified connectomes (as indicated by the legend) vs. fitted against the original connectome (see Methods). (F) In-degree distributions of rewired connectomes by layer. Same colors as in E, with black representing the original connectome.

### Rewiring of VIP+ Interneuron Connectivity in a Detailed Model of Rat Somatosensory Cortex

We employed the connectome manipulation framework to rewire interneuron connectivity in a detailed anatomical (Reimann et al., 2024) and physiological (Isbister et al., 2024) network model of the rat somatosensory cortex (Figure 4A; see details in Methods: *Detailed Network Model of the Rat Somatosensory Cortex*). We introduced a preference for VIP+ interneurons to target other inhibitory neurons with their connections (Pfeffer, Xue, He, Huang, & Scanziani, 2013; Pi et al., 2013) that was not present in the original network model (Reimann et al., 2024). The original (baseline) connectome of the network model is based on the detection of axo-dendritic appositions as potential synapses, followed by a target-unspecific pruning step that prefers multi-synaptic connections (Reimann et al., 2015). In order to introduce a preference for VIP+ interneurons to target other inhibitory neurons, we pruned the original (unpruned) set of appositions originating from VIP+ neurons (i.e., bitufted and small basket cells; m-types BTC and SBC) in layers 2/3, 4, 5, and 6 such that 96.5 % of their potential synapses on non-inhibitory target neurons were removed (Reimann et al., 2024), thereby reproducing inhibitory targeting trends found in MICrONS data (Schneider-Mizell et al., 2023). Based on this pruning rule, we extracted new adjacency and synaptome (i.e., numbers of synapses per connection) matrices for the connectivity within the central cortical column (Figure 4A, red hexagon), as well as the exact synapse positions on the dendrite, which were stored as generic models of types `LookupTableModel` (adjacency, synaptome) and `PropsTableModel` (positions) respectively (see Table S1).

We then rewired the connectivity originating from BTC and SBC interneurons in the central cortical column using the `conn rewiring` code module (see Table S3), while leaving all other connectivity unchanged. Physiological parameters and axonal delays of the new connections were drawn from stochastic models of types `Conn- PropsModel` and `LinDelayModel` respectively (see Table S1) that had been fitted against the original connectome beforehand (see Methods: *Fitting Physiological Parameter Models for VIP+ Pathways*). We employed the `LookupTableModel` storing the adjacency matrix as deterministic connectivity description, i.e., containing only connection probability values of zeros and ones as determined by the matrix. The numbers of synapses for each individual connections were taken from the `LookupTableModel` storing the synaptome matrix, and the exact synapse positions on the dendrites were externally loaded from the above-mentioned `PropsTableModel` (i.e., `external` option; see Table S4). We did not keep or reuse connections or their physiological parameterization in case they had already existed in the baseline connectome. Instead, pathway-specific physiological parameter values were independently drawn for new connections from the before-fitted connection properties model (i.e., `randomize` option; see Table S4). Likewise, synaptic delays were drawn from the before-fitted axonal delay model depending on the Euclidean distance between the pre-synaptic soma and the synapse position on the post- synaptic dendrite. The rewiring run was launched in parallel on five nodes of a computing cluster using 500 data splits.

Overall, the number of synapses remained relatively constant (*∼*25 % increase; Table 3), but they were spread over a much larger number of connections (over 700 % increase). Consequently, individual inhibitory connections were formed by a much lower number of synapses per connection (Figure 4B). Note that the low number of synapses per connection is indeed a feature of the electron-microscopic mouse dataset used in (Schneider-Mizell et al., 2023), while the baseline connectome matches the higher mean number of synapses per connection from paired light-microscopic reconstructions in rat (Reimann et al., 2015). We confirmed that synapses from BTC and SBC neurons were mainly targeting inhibitory (15-fold increase) instead of excitatory (two-fold decrease) neurons (Figure 4B, C; Figure S1). This is further validated by looking at the distribution of BTC and SBC synapses on the dendritic morphologies of two exemplary inhibitory and excitatory neurons before and after rewiring (Figure 4D; extensive list of examples in Figure S3 and S4 respectively). Importantly, physiological synapse parameters were found to be largely preserved after rewiring (Figure S2).

**Table 3.** Connection and synapse counts in rewired interneuron connectivity. The table contains the overall numbers of connections (#Conn) and synapses (#Syn) of the connectome with rewired interneuron connectivity with the respective differences to the original (baseline) connectome, in absolute terms (Diff) and percentages (%). Only the connectivity in the central cortical column originating from BTC/SBC source types which was subject to rewiring is considered here.

| Connectome | #Conn | Diff | % | #Syn | Diff | % |
| --- | --- | --- | --- | --- | --- | --- |
| **Baseline** | 90,200 | — | — | 1,276,686 | — | — |
| **Rewired** | 733,742 | 643,542 | 713.5 | 1,593,595 | 316,909 | 24.8 |

**Table 4.** Connection and synapse counts in rewired simplified connectomes. The table summarizes the numbers of connections (#Conn) and synapses (#Syn) of all simplified connectomes together with the respec- tive differences to the baseline connectome, in absolute terms (Diff) and percentages (%). Only the connectivity between excitatory neurons in the central cortical column of the network model which was subject to rewiring is considered here.

| Connectome | #Conn | Diff | % | #Syn | Diff | % |
| --- | --- | --- | --- | --- | --- | --- |
| **Baseline** | 7,203,362 | — | — | 31,212,240 | — | — |
| **1<sup>st</sup> order** | 7,205,703 | 2,341 | 0.032 | 26,603,307 | -4,608,933 | -14.766 |
| **2<sup>nd</sup> order** | 7,203,361 | -1 | -0.000 | 29,272,858 | -1,939,382 | -6.214 |
| **3<sup>rd</sup> order** | 7,203,364 | 2 | 0.000 | 30,173,482 | -1,038,758 | -3.328 |
| **4<sup>th</sup> order** | 7,203,362 | 0 | 0 | 30,332,403 | -879,837 | -2.819 |
| **5<sup>th</sup> order** | 7,203,362 | 0 | 0 | 31,153,461 | -58,779 | -0.188 |

We ran simulation experiments with both the original and the rewired connectome where we activated the rewired BTC and SBC interneurons by means of a constant 1 s current injection (Figure 4E; see details in Methods: *Current Injection Experiment*). During spontaneous activity before applying the current injection, we observed a slight increase in excitatory and slight decrease in inhibitory firing rates in the rewired connectome. During BTC and SBC activation, the firing rates of these types initially increased strongly, followed by a decay to a lower level that was still elevated compared to baseline conditions. The other inhibitory types (that were not injected with current) were largely inhibited by the newly created class of inhibitory targeting interneurons which was not the case in the original connectome. During this period, excitatory neuron firing rates decreased for both connectomes, but more strongly for the original connectome. This means that BTC/SBC interneurons still provided stronger inhibition than disinhibition to excitatory neurons. After the current injection ended, a brief rebound peak in the firing rates of the other inhibitory types was observed for the rewired connectome. The same result holds over a range of injected currents from 0.05 nA to 0.25 nA (Figure 4F).

### Simplified Connectivity of a Detailed Model of Rat Somatosensory Cortex

In a second experiment, we rewired the connectivity between excitatory neurons in the central column of the network model of the rat somatosensory cortex (Figure 4A; red hexagon). Rewiring was done by first simplifying the connectivity to one of the five following stochastic model descriptions (Gal et al. (2020); Figure 5A), i.e., by fitting their model parameters against the baseline connectome (see Methods: *Fitting Stochastic Models for Simplified Connectivity*), and then generating fixed instances of them, as illustrated in the “Rewire” case in Figure 1C (green) before.

**1**^**st**^ **order:** Constant connection probabilities between all pairs of neurons.

**2**^**nd**^ **order:** Distance-dependent connection probability between the pre- and post-synaptic neuron.

**3**^**rd**^ **order:** Bipolar distance-dependent connection probability based on two alternative distance-dependent probability functions for the pre-synaptic neuron being axially (i.e., along cortical depth axis) either above or below the post-synaptic neuron.

**4**^**th**^ **order:** Offset-dependent connection probability based on the axial and radial offsets between the pre- and post-synaptic neuron.

**5**^**th**^ **order:** Position-dependent connection probability based on the (absolute) axial position of the pre-synaptic neuron together with the axial and radial offsets between the pre- and post-synaptic neuron.

We rewired the whole connectivity between excitatory neurons within the central cortical column of the baseline connectome using the `conn rewiring` code module (see Table S3) based on the before-fitted stochastic 1^st^ to 5^th^ order connectivity models. For each of the connectivity models, we did this in an iterative way in order to match the total number of connections in the rewired connectomes to the baseline (see Methods: *Matching the Total Number of Connections in Rewired Connectomes*). Specifically, we ran between three and eight iterations in order to obtain close or exact matches of the overall numbers of connection, as summarized in Table 4. Again, each rewiring run was launched in parallel on five nodes of a computing cluster using 500 data splits.

Important aspects of connectivity, such as synapse locations on dendrites, pathway-specific physiological parameter distributions, and overall numbers of connections were preserved. Specifically, when realizing new con- nections trough individual synapses (i.e., synapse assignment and parameterization step, see Figure 1D2), we reused existing synapse positions on the dendrites (i.e., `reuse` option; see Table S4), but we did not keep or reuse in-degrees, the numbers of synapses per connection or their physiological parameterization. Instead, pathway-specific physiological parameter values were independently drawn from parameter distributions (i.e., `randomize` option; see Table 5B) given by a `ConnPropsModel` that had been fitted against the baseline connectome beforehand (see Methods: *Fitting Stochastic Models for Simplified Connectivity*). Likewise, synaptic delays were drawn from a `LinDelayModel`, that had been fitted beforehand, depending on the Euclidean distance between the new pre-synaptic soma and the newly assigned synapse position on the post-synaptic dendrite.

**Table 5.** Connection property distributions. Pathway-specific property distribution types that were fitted against the baseline con- nectome and used for drawing property values when realizing connections in (A) interneuron rewiring and (B) simplified connectomes.

| Property | Distribution type |  | Description |
| --- | --- | --- | --- |
|  | (A) Interneuron | (B) Simplified |  |
| n_syn_per_conn | — | discrete (integer) | Synapses per connection |
| conductance | gamma | gamma | Peak conductance (nS) |
| conductance_scale_factor | constant | constant | Conductance scaling factor |
| decay_time | truncated normal | truncated normal | Decay time constant (ms) |
| depression_time | gamma | gamma | Time constant for recovery from depression (ms) |
| facilitation_time | gamma | gamma | Time constant for recovery from facilitation (ms) |
| n_rrp_vesicles | discrete (integer) | discrete (integer) | Number of vesicles in readily releasable pool |
| syn_type_id | discrete (integer) | constant (integer) | Synapse type ID |
| u_syn | truncated normal | truncated normal | Utilization of synaptic efficacy |
| u_hill_coefficient | discrete | constant | Utilization scaling coefficient with calcium concentration |

We compared the structure of all rewired connectomes with the original (baseline) connectome (Figure 5B- D). As intended, only connections between excitatory neurons were rewired, by adding and deleting connections. The more extreme a simplification was, the fewer connections remained unchanged (Fig 5B, white). No connections from, to, or between inhibitory neurons were changed in any of the connectomes. The number of synapses forming a connection was drawn from pathway-specific distributions derived from the baseline connectome (see Figure S6). At the same time, while the overall number of connections was largely preserved (max. difference 0.032 %), connection counts in individual pathways could shift. Consequently, the total number of synapses could change quite drastically in the simpler connectomes (Figure 5C and Table 4): -15 % difference for the 1^st^ order connectome, decreasing to -0.2 % difference for the 5^th^ order connectome. Importantly, preserving the average number of synapses per connection of individual pathways rather than overall synapse count was necessary to preserve biologically parameterized amplitudes of post-synaptic potentials (PSPs). We further validated the simplifications on the level of layer-wise connection probabilities (Figure 5D). In the 1^st^ order connectome, we indeed found a uniform connection probability distribution between all layers which was very different from the baseline. The 2^nd^ order connectome already captured some of the structure present in the baseline; however, the layer-wise connectivity was completely symmetric and lacked any anisotropy. In the 3^rd^ order and higher connectomes, more and more of the underlying structure was captured, and the difference in connection probability in the 5^th^ order connectome was relatively small. This indicates that simplifying connectivity by taking positions and offsets into account closely approximates, on average, the underlying connectivity structure, in line with Gal et al. (2020). Also, we tested to what degree the simplifications of a given stochastic model order destroyed the structure that is captured by all other orders (Figure 5E; see Methods: *Model Order Validation of the Simplified Connectomes*). As expected, we found that a simplified connectome of a given order fully comprised all models of lower or equal order, i.e., an *n*^th^ order connectome is indistinguishable from the original connectome at all model orders below or equal *n*. Conversely, at higher model orders, substantial errors were visible. We further found that at the level of incoming connectivity the structural diversity is lessened in the simplified connectomes (Figure 5F). Finally, we also validated the mean numbers of synapses per connection and important physiological synapse parameters per pathway, i.e., for each pair of pre- and post-synaptic m-types (Figure S6). We found that the means and standard deviations of the pathway-specific parameter distributions from the baseline connectome were largely preserved in the rewired connectomes. Only for axonal delays, we found deviations in some of the pathways. This can be explained by the fact that unlike the other synapse parameters, axonal delays were not modeled in a pathway-specific way taking morphological differences into account, but by fitting the overall statistics of distance-dependent axonal delays.

We then ran a series of network simulations in order to quantify functional changes in the simplified connectomes. Specifically, we **recalibrated** the five rewired circuits to exhibit *in vivo*-like spontaneous activity using the calibration algorithm described in Isbister et al. (2024) (see details in Methods: *Recalibration of the Simplified Circuits*), in order to dissociate the effect of the redistribution of pathway strengths from the effect of higher-order structure. Briefly, the calibration iteratively parameterizes layer-specific conductance injections into the neurons until their firing rates match expected values. These injections represent the extrinsic inputs from regions that are not part of the network model. The expected values were a constant fraction of the firing rates observed in *in vivo* recordings. This fraction, *P*_*FR*_, was set to values below or equal to 1.0 to compensate for the presence of silent and hence “invisible” neurons (Buzsáki & Mizuseki, 2014; Olshausen & Field, 2006; Wohrer, Humphries, & Machens, 2013). We use the notation L*k*E and L*k*I to denote excitatory (E) and inhibitory (I) populations in layer *k* respectively. Without recalibration, the spontaneous activity for some simplified connectomes was in a synchronous activity state characterized by bursts of activity throughout all layers except layer 1 (at *P*_*FR*_ = 0.8; Figure 6A, left). This indicates that inhibition was no longer able to keep recurrent excitation under control. After several iterations of the calibration algorithm, the spontaneous activity was in an asynchronous activity state closely matching the expected firing rates, e.g., Figure 6A (right), for most values of *P*_*FR*_ (Figure 6B). For most simplified connectomes, only three iterations were required (Figure 6C) whereas the 2^nd^ order connectome (distance-dependent connectivity) had remaining errors even after five iterations (Figure 6B, red arrow; Figure S7; Figure S8). This highlights the importance of biologically realistic connectomes in order to obtain *in vivo*-like activity.

**Figure 6.**
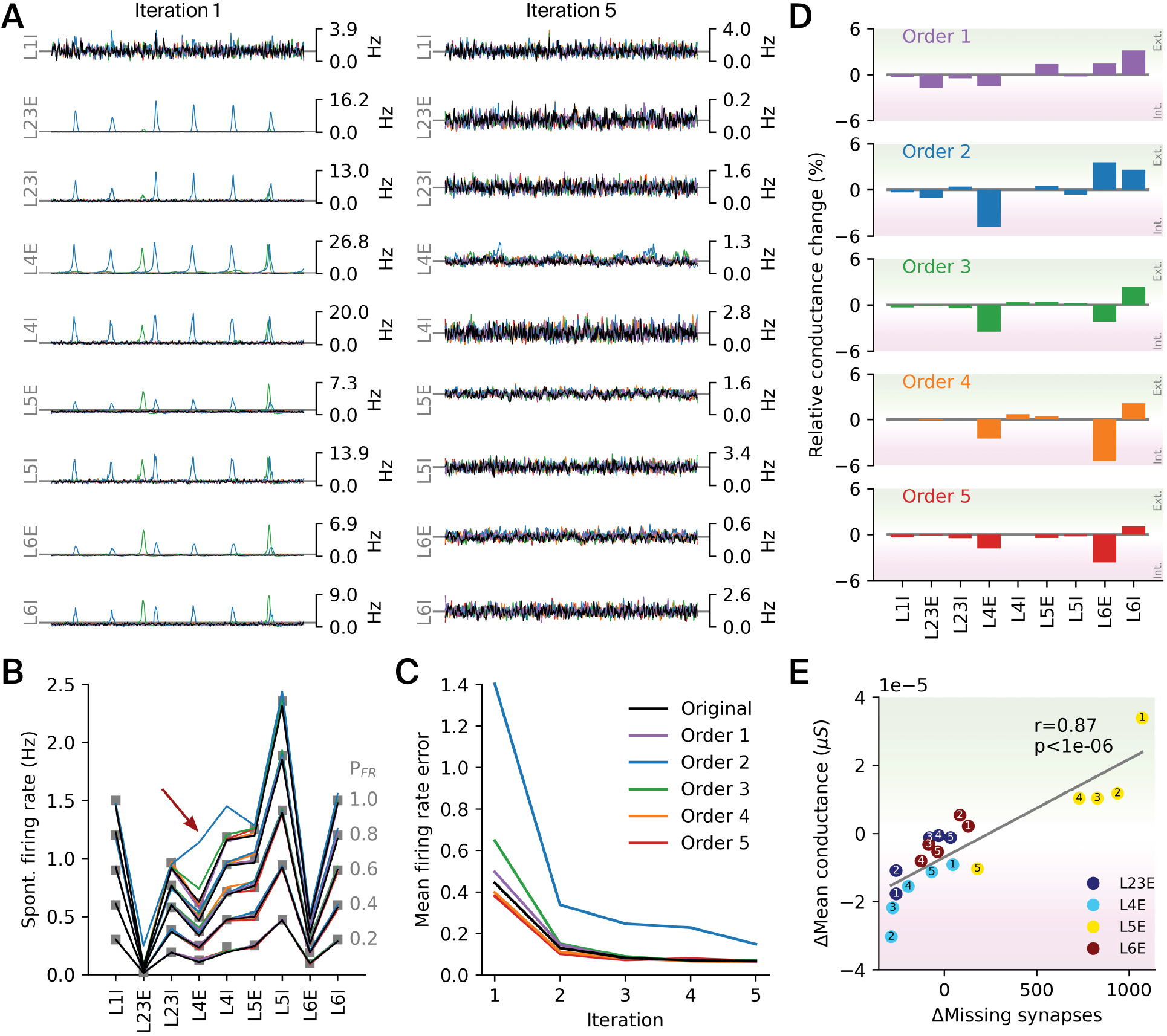
Functional implications of simplified connectivity. (A) Instantaneous firing rates of 5 s spontaneous activity of excitatory (E) and inhibitory (I) populations across layers for each of the rewired connectomes before (i.e., iteration 1) and after (i.e., iteration 5) recalibration, shown for an *in vivo* fraction *P*_*F R*_ = 0.8 (cf. Figure S8 with *P*_*F R*_ = 1.0). (B) Spontaneous firing rates after recalibration, closely matching *in vivo* references (grey squares) for most values of *P*_*F R*_ (see Methods), except for the 2^nd^ order connectome at *P*_*F R*_ = 1.0 (red arrow). (C) Mean firing rate error, computed as the Euclidean distance between the measured and the reference rates averaged over all *P*_*F R*_ values, for five calibration iterations. (D) Changes in mean conductance injection relative to baseline after recalibration, indicating neuron populations shifted towards a more externally (positive change) or internally (negative change) driven regime through rewiring. (E) Mean conductance injection versus mean number of missing excitatory synapses (both relative to baseline) for E populations across layers, as indicated by the legend. Small numbers denote the orders (1 to 5) of the rewired connectomes. The mean number of missing synapses was estimated assuming a reference density of 1.1 synapses/*μ;*m on the dendrites of individual neurons (Markram et al., 2015).

As the layer-specific injections represent extrinsic inputs, their changes during recalibration indicate to what degree a population is driven by intrinsic versus extrinsic populations (Figure 6D). We found that L4E neurons required less conductance injection compared to the baseline connectome, which implies that their activity was shifted towards a more internally driven spontaneous activity regime. This effect was most pronounced in the 2^nd^ order connectome and got successively weaker for more complex models. Moreover, we observed an inverse shift for L6I neurons towards a more externally driven regime, despite the fact that afferent connections to I populations had not been rewired. This effect was most pronounced in the 1^st^ order connectome. For L6E neurons, we observed diverse effects, being more externally driven in 1^st^ and 2^nd^ order, but more internally driven in 3^rd^ to 5^th^ order connectomes. This indicates the importance of the anisotropy of connectivity introduced in the 3^rd^ order connectome. As indicated before (Figure 5F), the incoming connectivity was redistributed, with some layers receiving more and some layers receiving less synaptic input in the simplified connectomes. As expected, we found that the additional amount of conductance required in all E populations was highly correlated with the change in afferent excitatory synapse count (Figure 6E; Pearson correlation coefficient *r* = 0.87; *p <* 10^*−*6^ based on a two-sided Wald test with t-distribution). This confirms our previous interpretation that the strength of conductance injection is an indication of how much a population is externally versus internally driven. Even though firing rates matched baseline after recalibration, the correlations of spiking activity were quite different in the simplified connectomes (Figure S9). While correlations in the baseline connectome steadily increased from superficial to deeper layers, the pattern was less clear cut in the simplified connectomes. In the least simplified 5^th^ order connectome correlations were most similar to baseline, but generally lower, especially in layer 6. This change is the result of reduced higher-order structure after the impact of a redistribution of pathway strengths has been controlled for.

## DISCUSSION

We present in this work a connectome manipulation framework that allows causal structure–function relationships to be studied in a systematic and reproducible way. We have demonstrated its utility in two exemplary applications using a detailed network model of the rat somatosensory cortex (Isbister et al., 2024; Reimann et al., 2024). In one experiment, we increased the inhibitory targeting specificity of VIP+ interneurons based on trends found in mouse EM data. In this case, we employed the framework to transplant specific connectivity provided by an adjacency matrix together with the numbers of synapses per connection in a deterministic way. In another experiment, we decreased the biological realism of the network model and studied the effect of such manipulation by rewiring the connectome based on simplified stochastic connectivity rules. In this case, we utilized the framework to rewire the entire connectivity between excitatory neurons in a stochastic way. Both of these experiment could supposedly have been conducted in an ad-hoc way without use of such framework. However, this would have required us to solve several challenges our framework inherently takes care of. Specifically, it allowed us to fit five simplified stochastic models of connectivity against the baseline connectome (Figure 5A; Figure S5). It enabled us to evaluate them during rewiring in order to generate connectome instances on the connection level (Figure 1D1). It ensured that the synaptic physiology of individual pathways was preserved, by fitting property distributions to the baseline connectome and evaluating them while turning the connectome into a description at the synapse level (Figure 1D2). It preserved the patterns of innervation of dendritic compartments by reusing existing synapse locations for the new connections (Table S5). While for stochastic models of connectivity the exact number of connections is in general not predetermined, the framework provided means to match the numbers to the baseline connectome (Table 4). It provided validations of the rewired connectomes on the connection and synapse level by structural comparison with the baseline (Figure 4B, C; Figure 5B-D; Figure S1; Figure S2; Figure S6). It offered high performance due to its parallel architecture (Figure 3), allowing us to rapidly rewire *∼*7M connections and *∼*30M synapses in less than 10 minutes on five computation nodes. Finally, it generated new SONATA circuits that could be readily used in simulation experiments (Figure 4E, F; Figure 6). Altogether, this shows that having such a framework as a reference tool greatly helps to standardize and reproduce *in silico* manipulation experiments.

Importantly, we have demonstrated that running connectome manipulations results in rewired connectomes as intended, i.e., as configured by the user. Whether this makes biological sense or not is up to the user to evaluate; the framework provides validation plots for this purpose. For example, the user could try to fit a distance-dependent connectivity description based on a parametric function (e.g., exponential, as in the 2^nd^ and 3^rd^ order models) but which does not reflect the underlying shape of distance-dependent connection probabilities extracted from data. In such a case, a refined parametric function or a non-parametric version of such a model could be introduced. We found for example that complex exponential functions provide a very good fit to the data (Figure S5). Also, the user could try to fit such a model to a connectome which does not have a well-defined notion of distance and/or axis alignment due to its complicated geometry, such as hippocampus (Romani et al., 2024) or other species like C. elegans (Witvliet et al., 2021) and insects (Winding et al., 2023). In such a case, a suitable position mapping to a coordinate systems which would take the curvature and/or laminar structure into account (Bolaños-Puchet et al., 2024) would be required. The same applies to long-range connectivity between different cortical regions in which case the source and target regions would have to be mapped to a common, local coordinate system first. In general, the higher the model order is, the closer the non-random expression patterns of triplet motifs and simplices found in experimental data can be reproduced, although not exactly (Egas Santander et al., 2024; Gal et al., 2020). A possible reason for this could be that these simplified stochastic models describe connection probabilities only on average, but don’t take the fine-grained structure of individual neurons into account. Also, in our present work and in Egas Santander et al. (2024), the models were fitted against the connectivity among excitatory neurons as a whole. Refinements could be made by fitting individual models for all pairs of layers and/or m-types, in order to better incorporate their individual geometrical characteristics. Aside from that, when fitting stochastic models to data, there is an inherent risk of overfitting. Intuitively, the risk increases the more parameters a model has (e.g., non-parametric 4^th^ and 5^th^ order models) and the sparser and/or noisier the data are. In order to reduce the risk of overfitting, parametric versions of these models could be added to the framework which would have a substantially lower number of parameters to fit. Moreover, the framework has a built-in functionality for cross-validation (see Section: *Model Fitting with Cross-Validation*) which can be utilized when fitting stochastic connectivity as well as synapse physiology models. However, synaptic variability may not be accurately captured by independent model distributions. For example, correlations between physiological parameters have been found before (Arel- lano, Benavides-Piccione, DeFelipe, & Yuste, 2007; Harris & Stevens, 1989; Schikorski & Stevens, 1999); we therefore implemented an option to specify correlations by means of a covariance matrix. Also, Ecker et al. (2024) predicted that central edges of a network are also physiologically stronger, a characteristic which is currently not captured by our stochastic models.

In our two exemplary experiments the framework allowed us to make several predictions. We predict that during spontaneous activity, activation of inhibitory targeting VIP+ interneurons reduces the firing rate of excitatory populations even though only 33 % of the synapses are targeting excitatory neurons (Figure 4). That is, although this group mostly shuts down firing of other interneurons the effect of this disinhibition is still weaker than its direct inhibition. Additionally, we learned that the removal of higher-order structure of connectivity when only the distance-dependent trends remained led to a transitions from an asynchronous to a synchronous state which had to be compensated for by reducing the amount of excitation from extrinsic sources (Figure 6). This was especially evident in layer 4, indicating that the role of layer 4 as input layer (Douglas & Martin, 2004; Miller, 2016) may be tied to the higher-order structure of recurrent connectivity. Structurally, the reduction or removal of higher-order structure led to a redistribution of strengths of layer-specific pathways. This is expected, as neuronal networks with more complex structure are associated with long-tailed degree distributions, while simplified networks have more homogeneous distributions. But it highlights the difficulty of dissociating first-order statistics, such as pathway strengths, from higher-order statistics in morphologically detailed models. Additionally, we found that in less realistic connectomes it may be harder to achieve a biologically realistic state (Figure S8), indicating the importance of properly modeling the higher-order connectivity structure. Finally, differences in the complexity of higher-order structure led to differences in the laminar patterns of spiking correlations.

Even though our two experiments were just brief outlines of full-scale experiments, we already gained interesting insights. In the future, such experiments could be done more extensively by direct use of EM reconstructions (MICrONS Consortium et al., 2021; Winding et al., 2023). For this to work, one would first need to create a SONATA version of an EM dataset, which involves modeling both the set of neurons and the connectivity between them. Then, one can study it by creating simplified but otherwise equivalent versions of the connectome as we have done before. This would provide insights into the importance of higher-order structure in a biologically measured and not merely predicted connectome. Also, the importance of individual pathways can in principle be studied by simply removing them, but such a severe manipulation may not be too insightful. A more fine-grained control for degrading them is provided by use of our manipulation framework. Recently, the higher-order network structure has been causally linked to important functional properties, such as reliability, efficiency, and population coupling (as defined by Okun et al. (2015)), using fine-grained and targeted manipulations implemented in our connectome manipulation framework (Egas Santander et al., 2024). We believe that following such ideas and going from purely correlational metrics to actual causation is a very promising way to develop the full potential of highly anticipated and valuable EM reconstructions. The need for investigating causal interaction in order to understand brain function has been pointed out before (Reid et al., 2019).

As our framework is inherently equipped to interpret connectivity at different levels of abstraction (Figure 1C), it can be used to bridge the scales between more and less detailed network models. For example, it could be employed to wire a detailed network model according to architectures used in machine learning in order to study under what conditions they would lead to biologically realistic results, thereby tightening the loop between neuroscience and artificial intelligence (Aru, Larkum, & Shine, 2023; Gershman, 2024; Gopinath, 2023; Surianarayanan, Lawrence, Chelliah, Prakash, & Hewage, 2023; Verzelli, Tchumatchenko, & Kotaleski, 2024). Also, by use of the open SONATA format (Dai et al., 2020), our framework supports not only biologically detailed but also point neuron network models, allowing for example the transformation of a connectome from a detailed into a point neuron network model. RÖssert et al. (2017) describes the adjustments to synaptic parameters required for such an endeavor that could be easily implemented as part of the operations described in Figure 1D2 in the context of the transplant functionality (Figure 1C, blue). Together with the recent publications of openly available large-scale models (Billeh et al., 2020; Dura-Bernal et al., 2023; Isbister et al., 2024), this makes our work also relevant for the point neuron model community.

Taken together, our framework serves as a flexible starting point for manipulating connectomes in a systematic and reproducible way, which can be easily extended and adapted to individual use cases (see *Extensions* in Table 1). New code modules can be simply integrated into the existing framework, such as new types of stochastic models, tools for fitting them against existing data, new manipulation operations, more specific synapse placement rules, and additional structural validations. Together with the ability to actually simulate such manipulated connectomes, this represents a powerful tool for fully understanding the role of connectivity in shaping network function.

## METHODS

### Details of the Connectome Manipulation Framework

#### Model Fitting with Cross-Validation

For fitting stochastic models to existing data, our framework optionally provides *k*-fold cross-validation (CV), which is a validation technique to prevent overfitting, in a semi-automated way. Stochastic models with a high number of parameters are in general more susceptible to overfitting. The number of model parameters are explicitly stated in Table S1. For example, the connection probability models of 1^st^ to 3^rd^ orders are parametric models with a relatively low number of parameters (only 1 in 1^st^ order, up to 10 in complex 3^rd^ order), as they are based on underlying mathematical functions. In contrast, the 4^th^ and 5^th^ order models are non-parametric in the sense that no underlying shape of the probability function is assumed, i.e., the number of model parameters is given by the specified number of data bins.

The user can specify the number of CV folds *k* (*k ≥* 2), either as a command line argument (see README file on GitHub) or in the model building configuration (see *Configuration file structure* in the Documentation). In so doing, the input data, i.e., the pre- and post-synaptic sets of neurons, will be randomly partitioned into *k* equal-sized folds, *k −* 1 of which will be used as a training set for fitting a model, and the remaining one as an unseen testing set which is plotted against the fitted model for validation purposes. In total, *k* different such models and validation plots will be generated, using each of the folds exactly once as a testing set. Based on these validation plots, the user can evaluate how well a fitted model generalizes to unseen data and how similar the *k* individual models are. In the end, the user could even create an combined model by averaging the respective parameter values of the *k* individual models. A minimal working CV example can be found in the `/examples` folder in the GitHub repository.

#### Matching the Total Number of Connections in Rewired Connectomes

When fitting stochastic connection probability models (as in Table S1) against existing connectomes and using them for rewiring, the resulting number of connections in the rewired connectome will in general not exactly match the number of connections in the original connectome for three reasons. First, stochastic connection probability models are in general simplified descriptions of connectivity which may not capture the underlying shape of the actual connection probabilities exactly (e.g., approximating distance-dependent connectivity by an exponential function). Second, stochastic connectivity models are not evaluated globally but locally for independently drawing incoming connections for each post-synaptic neuron (see first algorithmic step in Figure 1D1), allowing for efficient parallel processing (see Figure 3). Third, because of the stochastic nature of such probability models, the exact number of resulting connections varies in different random instances.

For applications where matched numbers of connections are desirable, our framework provides functionality for matching the overall number of connections as close as possible to the original number, based on the following assumptions. Rewiring is done by computing connection probabilities *p*_*ij*_ for all pre-synaptic neurons *i* to be connected to a given post-synaptic neuron *j* based on a stochastic model of connectivity. A specific instance (realization) randomly drawn from *p*_*ij*_ for each post-synaptic neuron *j* has *N*_*in,j*_ incoming connections. Thus, the expected number of incoming connections of a post-synaptic neuron *j* on average can be computed as

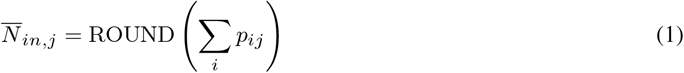

For adjusting the resulting number of connections, we implemented a global probability scaling factor *p*_*scale*_ into the rewiring operation which scales the connection probability 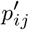 given by a stochastic connectivity model, i.e., 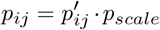 (by default, *p*_*scale*_ = 1.0). Using such global factor has the advantages that it neither changes the overall shape of the probability function nor introduces any dependencies between parallel processes or biases depending on the number of data splits.

In order to rapidly predict the total number of connections on average (i.e., independent of the random seed) when using a given connection probability model (incl. scaling factor *p*_*scale*_), we also implemented an “estimation run” option as part of the rewiring operation (see Table S4). This options allows a rewiring operation to be executed with early stopping, without generating an actual connectome or output file, but writing the average number of incoming connections *N*_*in,j*_ according to Eq. 1 into a data log file. The advantage of such an “estimation run” is that it facilitates obtaining the values *p*_*ij*_ as in Eq. 1 given all selected rewiring options and interdependencies that may exist, most importantly the selected pre- and post-synaptic neuron populations and the choice of the probability model which in turn may depend on an (optional) position mapping model. Using these values from the data log, the total number of connections can be computed as sum over all post-synaptic neurons *j* that are subject to rewiring, i.e.,

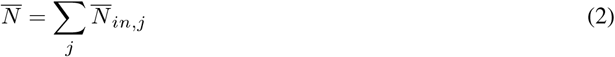

Based on this number, a scaling factor *p*_*scale*_ can be computed by

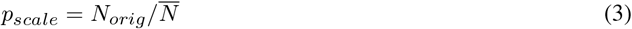

where *N*_*orig*_ is the total number of connections in the original connectome (for the same selection of pre- and post-synaptic populations).

While the above functionality allows the adjustment of the resulting numbers on average, a single random instance may still deviate. Therefore, we implemented yet another option for optimizing the drawn number of connections in a single instance. Specifically, the number of incoming connections for each post-neuron will be optimized to match its average number of connections. This is done by repeating the random generation up to 1,000 times and keeping the instance with the number of connections exactly or as close as possible matched to the average.

So, for closely matching the overall number of connections, we propose the following semi-automatic two-step procedure:

Step 1: Matching the mean

- Run estimation runs iteratively, until the total predicted number of connections *N* in the rewired connectome (Eq. 2) exactly matches the total number of connections *N*_*orig*_ in original connectome. A few iterations are usually required since a non-linear rounding operation to integer numbers of connections is involved (see Eq. 1).
- After each iteration, compute a new scaling factor *p*_*scale,new*_ using Eq. 3.
- Update *p*_*scale*_ = *p*_*scale*_ *· p*_*scale,new*_ based on this new estimate, to be used in the next iteration.
- Convergence is reached if *p*_*scale,new*_ = 1.0, i.e., the expected number of connections exactly matches the number in the original connectome. However, convergence is not guaranteed and values may oscillate around the theoretical optimum 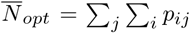 (i.e., without rounding). In such case, the value closest to the optimum should be used.

Step 2: Matching an instance to the mean

- Use converged (or closest) value *p*_*scale*_ from step 1.
- Enable option to optimize numbers of connections in a single instance.
- Run actual rewiring operation for a specific random seed (i.e., no estimation run).

Even though this procedure usually finds a close solution after only a few iterations, it is not guaranteed to converge to the exact number under all circumstances. This is mainly due to the discrete nature of the local rewiring where integer numbers of incoming connections are independently drawn for each post-synaptic neuron, and the parallel processing architecture which limits the exchange of information, such as the actually drawn numbers of connections, among independently processed data splits.

### Detailed Network Model of the Rat Somatosensory Cortex

We employed connectome manipulations in a detailed anatomical (Reimann et al., 2024) and physiological (Isbister et al., 2024) network model of the non-barrel rat somatosensory cortex (Figure 4A). While the full model contains over 4M neurons, we utilized the openly available 1.5 mm diameter subvolume composed of seven hexagonal columns, which has been released under the Digital Object Identifier (DOI) 10.5281/zenodo.8026353. This data-driven subvolume model consists of over 210k biophysically detailed neurons belonging to 60 different morphological types (m-types) and 208 different morpho-electrical types, which are connected by over 400M synapses with probabilistic transmitter release and 5 distinct forms of short-term dynamics of depression and facilitation. The connectome is based on axo-dendritic appositions, which has been demonstrated to reproduce non-random trends identified in biological networks (Gal et al., 2017). As depicted in Figure 4A, we applied connectome manipulations only to connections between 30,190 neurons within the central cortical column of the seven column subvolume, in order to avoid potential edge effects. For all manipulations with geometry-dependent connection probabilities (i.e., 2^nd^ to 5^th^ order), we used the flat coordinate system mapping released in Bolaños-Puchet et al. (2024) under DOI 10.5281/zenodo.10686776, with the x/y-axes parallel to the cortical layers, and the z-axis along the cortical depth.

For simulating the model, we used the CoreNEURON simulator (Kumbhar et al., 2019) together with the openly-available Neurodamus simulator control application (see Section: *Software and Data Availability*). As for manipulations, we only simulated neurons within the central cortical column of the seven column subvolume, using connections from the baseline or one of the manipulated connectomes between them. The neurons themselves were not subject to any manipulations and remained identical in all simulations.

In order to obtain *in vivo*-like spontaneous activity during simulations, we compensated for missing excitatory inputs that were external to the network model, as described in Isbister et al. (2024). Input compensation was given by statistically independent, population-specific somatic conductance injections from Ornstein-Uhlenbeck (OU) processes that would mimic aggregated random background synaptic inputs. The compensation mechanism was based on three meta-parameters: the extracellular calcium concentration *Ca*, the fixed ratio between standard deviation and mean of the underlying OU processes *R*_*OU*_, and a constant fraction *P*_*FR*_ of the population-specific *in vivo* reference values (De Kock, Bruno, Spors, & Sakmann, 2007; Reyes-Puerta, Sun, Kim, Kilb, & Luhmann, 2015), taking into account that extracellularly recorded firing rates are known to be overestimated to an unknown degree (Buzsáki & Mizuseki, 2014; Olshausen & Field, 2006; Wohrer et al., 2013). Unless noted otherwise, we used the calibration of OU-parameters from Isbister et al. (2024) on the full seven column subvolume for *Ca* of 1.05 mM, a ratio *R*_*OU*_ of 0.4, and an *in vivo* proportion *P*_*FR*_ of 0.3.

### Details of Rewired VIP+ Interneuron Connectivity

#### Fitting Physiological Parameter Models for VIP+ Pathways

For VIP+ interneuron rewiring, two physiological model descriptions were required: a stochastic model for realizing new connections by forming and parameterizing synapses, and a stochastic model for assigning their axonal delays. For realizing connections, a connection properties model of type `ConnPropsModel` (see Table S1) was fitted against the central column (Figure 4A, red hexagon) of the baseline connectome using the `conn props` code module (see Table S2), by extracting pathway-specific parameter distributions, i.e., for all 8 *×* 60 pairs of pre- and post-synaptic m-types that were subject of rewiring, of the distribution types as summarized in Table 5A. The numbers of synapses per connections were not drawn from that model, but provided deterministically through the synaptome. All remaining properties not listed in this table were not relevant for simulations and were set to zero. Pathways with less than 10 connections were treated as missing values and therefore gradually interpolated from similar pathways (as detailed in Table S2). For pathways with more than 10k connections, a random subset of 10k connections was used for distribution fitting.

A model for assigning linearly distance-dependent axonal delays of type `LinDelayModel` (see Table S1) was fitted against connections between the relevant pre- and post-synaptic m-types in the central column of the baseline connectome using the `delay` code module (see Table S2). We utilized a distance bin size of 50 *μ;*m and did not distinguish between individual pathways.

#### Current Injection Experiment

We ran simulation experiments of the central cortical column of both the baseline and the rewired connectome during which we activated the BTC and SBC interneurons by injecting a constant current. In each simulations, we injected one of the five current strengths 0.05 nA, 0.1 nA, 0.15 nA, 0.2 nA, and 0.25 nA respectively. The total simulation duration of 10 s was divided into four time windows in which the network activity was analyzed afterwards:

***W***_**1**,***Spont***_ … Spontaneous activity, from *t* = 2 s to 5 s

***W***_**2**,***Inj***_ … Current injection time window, from *t* = 5 s to 6 s

***W***_**3**,***Rec***_ … Recovery time window after injection, from *t* = 6 s to 7 s

***W***_**4**,***Spont***_ … Spontaneous activity, from *t* = 7 s to 10 s

For analyzing the activity, we computed the average firing rates *R* over three distinct populations of neurons:

***R***_***E***_ … Excitatory neurons

***R***_***I***,***Inj***_ … Injected inhibitory neurons (BTC and SBC m-types)

***R***_***I***,***\Inj***_ … Non-injected inhibitory neurons (i.e., all remaining inhibitory m-types)

Instantaneous population firing rates as shown in Figure 4E were estimated with a bin size of 10 ms and smoothed with a Gaussian kernel with a standard deviation of 1.0. Significant differences between baseline and rewired activity were computed as the negative decimal logarithm of the p-values obtained by a Wilcoxon rank-sum test applied on 200 ms sliding windows of the instantaneous firing rates. Also, average firing rates of all time windows and populations were computed for all current strengths, results of which can be seen in Figure 4F.

### Details of Simplified Connectomes

#### Fitting Stochastic Models for Simplified Connectivity

We fitted five simplified stochastic connection probability models from 1^st^ to 5^th^ order (Gal et al., 2020) against the connectivity between excitatory neurons in the central cortical column of the baseline connectome using the `conn prob` code module (see Table S2). The resulting 1^st^ order model was of type `ConnProb1stOrderModel` and just contained the average connection probability extracted from the data (*p*_*const*_ = 0.010; see Table S1). As all higher-order models depend on geometry, they required correct alignment of the coordinate axis with the cortical layers. We therefore employed a coordinate transformation of the neuron positions to a flat coordinate system, by linearly interpolating the voxel-based flat and depth coordinates of the network model (see Section: *Software and Data Availability*) using the `pos mapping` code module (see Table S2). We scaled the x/y-axis by a factor of 34.0 *·* 189.0 from normalized units to *μ;*m, and the z-axis by a factor of -1.0, i.e., along the negative cortical depth (in *μ;*m). The resulting position mapping extension of type `PosMapModel` (see Table S1) was then applied when fitting connectivity models and using them in manipulations.

For fitting the 2^nd^ and 3^rd^ order models, we first extracted binned connection probability values with a distance bin size of 50 *μ;*m, and then fitted complex exponential functions using the Python function scipy.optimize.curve fit to these probability values (see Table S2). The resulting connectivity models were of types `ConnProb2ndOrder- ComplexExpModel` (with *α*_*p*_ = 0.084, *β*_*p*_ = 0.000186, *γ* = 1.735, *α*_*d*_ = 0.017, *β*_*d*_ = 0.002) and `ConnProb- 3rdOrderComplexExpModel` (with *α*_*p−*_ = 0.087, *β*_*p−*_ = 0.000042, *γ*_*−*_ = 2.0, *α*_*d−*_ = 0.024, *β*_*d−*_ = 0.001, *α*_*p*+_ = 0.081, *β*_*p*+_ = 0.001004, *γ*_+_ = 1.444, *α*_*d*+_ = 0.013, *β*_*d*+_ = 0.003) respectively (see Table S1; Figure S5). Unlike the simplified models proposed in Gal et al. (2020), we used reduced, radial symmetric versions of the 4^th^ and 5^th^ order models which had a radial component within in the x/y-plane and an axial component along the z-axis, and position-dependence in the 5^th^ order model only along the z-axis. For constructing these models, we first extracted binned connection probability values using a radial offset binning from 0 to 450 *μ;*m and an axial offset binning from -1550 to 650 *μ;*m in steps of 50 *μ;*m, and for the 5^th^ order model in addition a position binning from -2400 to 200 *μ;*m in steps of 200 *μ;*m. The resulting connectivity models were then based on bi- and tri-linear interpolation given these binned probability values on the regular grids of data bins using the Python function scipy.interpolate.interpn, and were of types `ConnProb4thOrderLinInterpnReduced- Model` and `ConnProb5thOrderLinInterpnReducedModel` respectively (see Table S1). We excluded bins with less than 100 data points in order to reduce noise in all model fits.

Again, stochastic models for parameterizing new connections and assigning axonal delays were required. We used the same types of models, a `ConnPropsModel` and a `LinDelayModel`, as for interneuron rewiring (see Section: *Fitting Physiological Parameter Models for VIP+ Pathways*) but which were fitted against the connectivity between 18 *×* 18 m-types of excitatory neurons in the central cortical column here. The property distribution types as summarized in Table 5B were used, including a distribution for numbers of synapses per connection. All other model fitting parameters were kept the same as for VIP+ interneuron rewiring.

#### Model Order Validation of the Simplified Connectomes

For validating the model order of the rewired simplified connectomes, we refitted each of the stochastic 1^st^ to 5^th^ order models against each of the five simplified connectomes, resulting in 25 model fits whose connection probabilities were given by 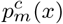 for model order *m* and simplified connectome order *c*; *x* denotes the set of respective input parameters an order-*m* model depends on, i.e., no input for 1^st^ order, distance variable for 2^nd^ order, etc. For each model order *m*, we probed the corresponding probability function in steps of 10 *μ;*m of their respective input variables (i.e., distance, offset, position, etc.) and computed the mean-squared error (MSE) with respect to the probabilities 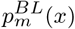 probed at the same input values of the model fits to the baseline connectome, i.e., the ones that had been used for rewiring in the first place. The MSE for a simplified connectome with order *c* was computed as

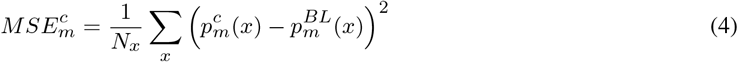

with *N*_*x*_ being the total number of probed input values of *x* for a given model order *m*. Specifically, we probed the probability functions at input values as follows: For *m* = 1, we used *N*_*x*_ = 1 since 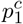 is a constant value without any dependencies; for *m* = 2, we used *N*_*x*_ = 251 distance values from 0 to 2500 *μ;*m; for *m* = 3, we used *N*_*x*_ = 501 (bipolar) distance values from -2500 to 2500 *μ;*m; for *m* = 4, we used *N*_*x*_ = 51 *·* 221 as given by 51 radial offset values from 0 to 500 *μ;*m and 221 axial offset values from -1600 to 600 *μ;*m; for *m* = 5, we used *N*_*x*_ = 51 *·* 221 *·* 201 as given by the same offsets as before and in addition 201 axial position values from -2000 to 0 *μ;*m. The resulting MSEs for all model orders and connectomes are shown in Figure 5E.

#### Recalibration of the Simplified Circuits

We employed the iterative calibration algorithm developed in Isbister et al. (2024) in order to calibrate population-specific OU-parameters (see Section: *Detailed Network Model of the Rat Somatosensory Cortex*) in the five circuits with rewired simplified connectomes so that their activity within the central cortical column would exhibit *in vivo*-like spontaneous activity. Likewise, we also recalibrated the original circuit with baseline connectome which had an initial calibration on the full seven column subvolume. Specifically, for each of the circuits, we ran five calibration iterations starting from the same initial calibration as the original circuit and using a calcium level *Ca* of 1.05 mM, a ratio *R*_*OU*_ of 0.4, and an *in vivo* proportion *P*_*FR*_ from 0.1 to 1.0 in steps of 0.1. After recalibration, we quantified the mean conductance injections required in the different excitatory and inhibitory populations and layers relative to the original circuit, results of which can be found in Figure 6D.

### Software and Data Availability

All software and data related to this article are openly available as follows:

#### Connectome-Manipulator

Python framework for connectome manipulations presented in this article, available under https://github.com/BlueBrain/connectome-manipulator, together with additional use case examples under `/examples`.

#### Connectome-Manipulator documentation

Documentation of the connectome manipulation framework, avail- able under https://connectome-manipulator.readthedocs.io. The documentation contains API references for model fitting, connectome manipulation, and structural comparison functions, as well as a reference page about their respective configuration file structures.

#### parquet-converters

External dependency required for automatically converting the individual output .parquet files produced by Connectome-Manipulator to an output connectome in SONATA format, available under https://github.com/BlueBrain/parquet-converters.

#### SSCx network model

Seven column subvolume of a detailed model of the rat somatosensory cortex (Isbister et al., 2024; Reimann et al., 2024) in SONATA format used as a basis for all manipulations in this article, available under DOI 10.5281/zenodo.8026353.

#### SSCx flat coordinates

Coordinate mapping for the SSCx network model to a flat coordinate system aligned with the cortical layers, which has been released in Bolaños-Puchet et al. (2024) under DOI 10.5281/zenodo.10686776.

#### SSCx connectome manipulation code

Repository with code and configuration files for applying manipulations to the SSCx network model, analyzing results, and reproducing the figures in this article, available under https://github.com/BlueBrain/sscx-connectome-manipulations.

#### SSCx connectome manipulation data

Dataset containing the resulting data of the manipulated SSCx network model, such as fitted stochastic models, manipulated connectomes, structural validations, as well as simulation data and analysis results, available under DOI 10.5281/zenodo.11402578.

#### Simulator software

Simulator software Neurodamus, which is a simulation control application for the NEURON simulator, released in Isbister et al. (2024) under DOI 10.5281/zenodo.8075202 for simulating the SSCx network model.

## AUTHOR CONTRIBUTIONS

- Conceptualization: C.P., M.W.R.
- Data curation: C.P.
- Formal analysis: C.P.
- Investigation: C.P., M.W.
- Methodology: C.P., J.B.I., K.K., M.W.R.
- Project administration: M.W.R.
- Software: C.P., O.A., K.K., M.W.
- Supervision: M.W.R.
- Validation: C.P., K.K.
- Visualization: C.P.
- Writing - original draft: C.P., M.W.R.
- Writing - review & editing: C.P., O.A., J.B.I., K.K., M.W., M.W.R.

## Supporting information

List of technical terms

## ACKNOWLEDGMENTS

This study was supported by funding to the Blue Brain Project, a research center of the É cole polytechnique fédérale de Lausanne (EPFL), from the Swiss government’s ETH Board of the Swiss Federal Institutes of Technology.

## SUPPLEMENTARY MATERIALS

Supplementary materials are available in a separate .pdf file.

## REFERENCES

Arellano, J. I., Benavides-Piccione, R., DeFelipe, J., & Yuste, R. (2007). Ultrastructure of dendritic spines: Correlation between synaptic and spine morphologies. Frontiers in Neuroscience, 1, 42. doi: 10.3389/neuro.01.1.1.010.2007

Aru, J., Larkum, M. E., & Shine, J. M. (2023). The feasibility of artificial consciousness through the lens of neuroscience. Trends in Neurosciences, 46(12), 1008–1017. doi: 10.1016/j.tins.2023.09.009

Azevedo Carvalho, N., Contassot-Vivier, S., Buhry, L., & Martinez, D. (2020). Simulation of large scale neural models with event-driven connectivity generation. Frontiers in Neuroinformatics, 14, 522000. doi: 10.3389/fninf.2020.522000

Bachmann, P. (1894). Zahlentheorie: Die analytische Zahlentheorie (Vol. 2). B.G. Teubner, Leipzig.

Billeh, Y. N., Cai, B., Gratiy, S. L., Dai, K., Iyer, R., Gouwens, N. W., … Arkhipov, A. (2020). Systematic integration of structural and functional data into multi-scale models of mouse primary visual cortex. Neuron, 106(3), 388–403. doi: 10.1016/j.neuron.2020.01.040

Bolaños-Puchet, S., Teska, A., Hernando, J. B., Lu, H., Romani, A., Schürmann, F., & Reimann, M. W. (2024). Enhancement of brain atlases with laminar coordinate systems: Flatmaps and barrel column annotations. Imaging Neuroscience, 2, 1–20. doi: 10.1162/imaga00209

Buzsáki, G., & Mizuseki, K. (2014). The log-dynamic brain: How skewed distributions affect network operations. Nature Reviews Neuroscience, 15(4), 264–278. doi: 10.1038/nrn3687

Dai, K., Hernando, J., Billeh, Y. N., Gratiy, S. L., Planas, J., Davison, A. P., … Arkhipov, A. (2020). The SONATA data format for efficient description of large-scale network models. PLOS Computational Biology, 16(2), e1007696. doi: 10.1371/journal.pcbi.1007696

Deco, G., & Hugues, E. (2012). Neural network mechanisms underlying stimulus driven variability reduction. PLOS Computational Biology, 8(3), e1002395. doi: 10.1371/journal.pcbi.1002395

De Kock, C., Bruno, R. M., Spors, H., & Sakmann, B. (2007). Layer-and cell-type-specific suprathreshold stimulus representation in rat primary somatosensory cortex. Journal of Physiology, 581(1), 139–154. doi: 10.1113/jphysiol.2006.124321

Douglas, R. J., & Martin, K. A. (2004). Neuronal circuits of the neocortex. Annual Review of Neuroscience, 27, 419–451. doi: 10.1146/annurev.neuro.27.070203.144152

Dura-Bernal, S., Griffith, E. Y., Barczak, A., O’Connell, M. N., McGinnis, T., Moreira, J. V., … Neymotin, S. A. (2023). Data-driven multiscale model of macaque auditory thalamocortical circuits reproduces in vivo dynamics. Cell Reports, 42(11), 113378. doi: 10.1016/j.celrep.2023.113378

Ecker, A., Egas Santander, D., Abdellah, M., Alonso, J. B., Bolaños-Puchet, S., Chindemi, G., … Reimann, M. W. (2024). Assemblies, synapse clustering and network topology interact with plasticity to explain structure-function relationships of the cortical connectome. eLife. doi: 10.7554/elife.101850.1

Egas Santander, D., Pokorny, C., Ecker, A., Lazovskis, J., Santoro, M., Smith, J. P., … Reimann, M. W. (2024). Heterogeneous and higher-order cortical connectivity undergirds efficient, robust and reliable neural codes. bioRxiv. doi: 10.1101/2024.03.15.585196

Gal, E., London, M., Globerson, A., Ramaswamy, S., Reimann, M. W., Muller, E., … Segev, I. (2017). Rich cell-type-specific network topology in neocortical microcircuitry. Nature Neuroscience, 20(7), 1004–1013. doi: 10.1038/nn.4576

Gal, E., Perin, R., Markram, H., London, M., & Segev, I. (2020). Neuron geometry underlies universal network features in cortical microcircuits. bioRxiv. doi: 10.1101/656058

Gershman, S. J. (2024). What have we learned about artificial intelligence from studying the brain? Biological Cybernetics, 1–5. doi: 10.1007/s00422-024-00983-2

Gopinath, N. (2023). Artificial intelligence and neuroscience: An update on fascinating relationships. Process Biochemistry, 125, 113–120. doi: 10.1016/j.procbio.2022.12.011

Harris, K. M., & Stevens, J. K. (1989). Dendritic spines of CA 1 pyramidal cells in the rat hippocampus: Serial electron microscopy with reference to their biophysical characteristics. Journal of Neuroscience, 9(8), 2982–2997. doi: 10.1523/jneurosci.09-08-02982.1989

Isbister, J. B., Ecker, A., Pokorny, C., Bolaños-Puchet, S., Santander, D. E., Arnaudon, A., … Reimann, M. W. (2024). Modeling and simulation of neocortical micro- and mesocircuitry. Part II: Physiology and experimentation. eLife. doi: 10.7554/elife.99693.1

Kumbhar, P., Hines, M., Fouriaux, J., Ovcharenko, A., King, J., Delalondre, F., & Schürmann, F. (2019). CoreNEURON: An optimized compute engine for the NEURON simulator. Frontiers in Neuroinformatics, 13, 63. doi: 10.3389/fninf.2019.00063

Lagzi, F., & Rotter, S. (2015). Dynamics of competition between subnetworks of spiking neuronal networks in the balanced state. PLOS ONE, 10(9), e0138947. doi: 10.1371/journal.pone.0138947

Litwin-Kumar, A., & Doiron, B. (2012). Slow dynamics and high variability in balanced cortical networks with clustered connections. Nature Neuroscience, 15(11), 1498–1505. doi: 10.1038/nn.3220

Markram, H., Muller, E., Ramaswamy, S., Reimann, M. W., Abdellah, M., Sanchez, C. A., … Schürmann, F. (2015). Reconstruction and simulation of neocortical microcircuitry. Cell, 163(2), 456–492. doi: 10.1016/j.cell.2015.09.029

MICrONS Consortium, Bae, J. A., Baptiste, M., Bishop, C. A., Bodor, A. L., Brittain, D., … Zhang, C. (2021). Functional connectomics spanning multiple areas of mouse visual cortex. bioRxiv. doi: 10.1101/2021.07.28.454025

Miller, K. D. (2016). Canonical computations of cerebral cortex. Current Opinion in Neurobiology, 37, 75–84. doi: 10.1016/j.conb.2016.01.008

Okun, M., Steinmetz, N. A., Cossell, L., Iacaruso, M. F., Ko, H., Barthó, P., … Harris, K. D. (2015). Diverse coupling of neurons to populations in sensory cortex. Nature, 521(7553), 511–515. doi: 10.1038/nature14273

Olshausen, B. A., & Field, D. J. (2006). What is the other 85 percent of V1 doing? In J. L. van Hemmen & T. J. Sejnowski (Eds.), 23 Problems in Systems Neuroscience (pp. 182–211). Oxford University Press. doi: 10.1093/acprof:oso/9780195148220.003.0010

Perin, R., Berger, T. K., & Markram, H. (2011). A synaptic organizing principle for cortical neuronal groups. Proceedings of the National Academy of Sciences, 108(13), 5419–5424. doi: 10.1073/pnas.1016051108

Pfeffer, C. K., Xue, M., He, M., Huang, Z. J., & Scanziani, M. (2013). Inhibition of inhibition in visual cortex: The logic of connections between molecularly distinct interneurons. Nature Neuroscience, 16(8), 1068–1076. doi: 10.1038/nn.3446

Pi, H.-J., Hangya, B., Kvitsiani, D., Sanders, J. I., Huang, Z. J., & Kepecs, A. (2013). Cortical interneurons that specialize in disinhibitory control. Nature, 503(7477), 521–524. doi: 10.1038/nature12676

Reid, A. T., Headley, D. B., Mill, R. D., Sanchez-Romero, R., Uddin, L. Q., Marinazzo, D., … Cole, M. W. (2019). Advancing functional connectivity research from association to causation. Nature Neuroscience, 22(11), 1751–1760. doi: 10.1038/s41593-019-0510-4

Reimann, M. W., Bolaños-Puchet, S., Courcol, J.-D., Egas Santander, D., Arnaudon, A., Coste, B., … Ramaswamy, S. (2024). Modeling and simulation of neocortical micro- and mesocircuitry. Part I: Anatomy. eLife. doi: 10.7554/elife.99688.1

Reimann, M. W., King, J. G., Muller, E. B., Ramaswamy, S., & Markram, H. (2015). An algorithm to predict the connectome of neural microcircuits. Frontiers in Computational Neuroscience, 9, 120. doi: 10.3389/fncom.2015.00120

Renart, A., Moreno-Bote, R., Wang, X.-J., & Parga, N. (2007). Mean-driven and fluctuation-driven persistent activity in recurrent networks. Neural Computation, 19(1), 1–46. doi: 10.1162/neco.2007.19.1.1

Reyes-Puerta, V., Sun, J.-J., Kim, S., Kilb, W., & Luhmann, H. J. (2015). Laminar and columnar structure of sensory-evoked multineuronal spike sequences in adult rat barrel cortex in vivo. Cerebral Cortex, 25(8), 2001–2021. doi: 10.1093/cercor/bhu007

Romani, A., Antonietti, A., Bella, D., Budd, J., Giacalone, E., Kurban, K., … Markram, H. (2024). Community-based reconstruction and simulation of a full-scale model of the rat hippocampus CA1 region. PLOS Biology, 22(11), 1–61. doi: 10.1371/journal.pbio.3002861

Rost, T., Deger, M., & Nawrot, M. P. (2018). Winnerless competition in clustered balanced networks: Inhibitory assemblies do the trick. Biological Cybernetics, 112(1-2), 81–98. doi: 10.1007/s00422-017-0737-7

Rössert, C., Pozzorini, C., Chindemi, G., Davison, A. P., Eroe, C., King, J., … Muller, E. (2017). Automated point-neuron simplification of data-driven microcircuit models. arXiv. doi: 10.48550/arXiv.1604.00087

Schikorski, T., & Stevens, C. (1999). Quantitative fine-structural analysis of olfactory cortical synapses. Proceedings of the National Academy of Sciences, 96(7), 4107–4112. doi: 10.1073/pnas.96.7.4107

Schneider-Mizell, C. M., Bodor, A., Brittain, D., Buchanan, J., Bumbarger, D. J., Elabbady, L., … Da Costa, N. M. (2023). Cell-type-specific inhibitory circuitry from a connectomic census of mouse visual cortex. bioRxiv. doi: 10.1101/2023.01.23.525290

Song, S., Sjöström, P. J., Reigl, M., Nelson, S., & Chklovskii, D. B. (2005). Highly nonrandom features of synaptic connectivity in local cortical circuits. PLOS Biology, 3(3), e68. doi: 10.1371/journal.pbio.0030068

Strata, P., & Harvey, R. (1999). Dale’s principle. Brain Research Bulletin, 50(5), 349–350. doi: 10.1016/S0361-9230(99)00100-8

Surianarayanan, C., Lawrence, J. J., Chelliah, P. R., Prakash, E., & Hewage, C. (2023). Convergence of artificial intelligence and neuroscience towards the diagnosis of neurological disorders—A scoping review. Sensors, 23(6), 3062. doi: 10.3390/s23063062

Verzelli, P., Tchumatchenko, T., & Kotaleski, J. H. (2024). Editorial overview: Computational neuroscience as a bridge between artificial intelligence, modeling and data. Current Opinion in Neurobiology, 84, 102835. doi: 10.1016/j.conb.2023.102835

Winding, M., Pedigo, B. D., Barnes, C. L., Patsolic, H. G., Park, Y., Kazimiers, T., … Zlatic, M. (2023). The connectome of an insect brain. Science, 379(6636), eadd9330. doi: 10.1126/science.add9330

Witvliet, D., Mulcahy, B., Mitchell, J. K., Meirovitch, Y., Berger, D. R., Wu, Y., … Zhen, M. (2021). Connectomes across development reveal principles of brain maturation. Nature, 596(7871), 257–261. doi: 10.1038/s41586-021-03778-8

Wohrer, A., Humphries, M. D., & Machens, C. K. (2013). Population-wide distributions of neural activity during perceptual decision-making. Progress in Neurobiology, 103, 156–193. doi: 10.1016/j.pneurobio.2012.09.004

