## Supplementary material for "A connectome manipulation framework for the systematic and reproducible study of structure–function relationships through simulations"

#### Supplementary Figures

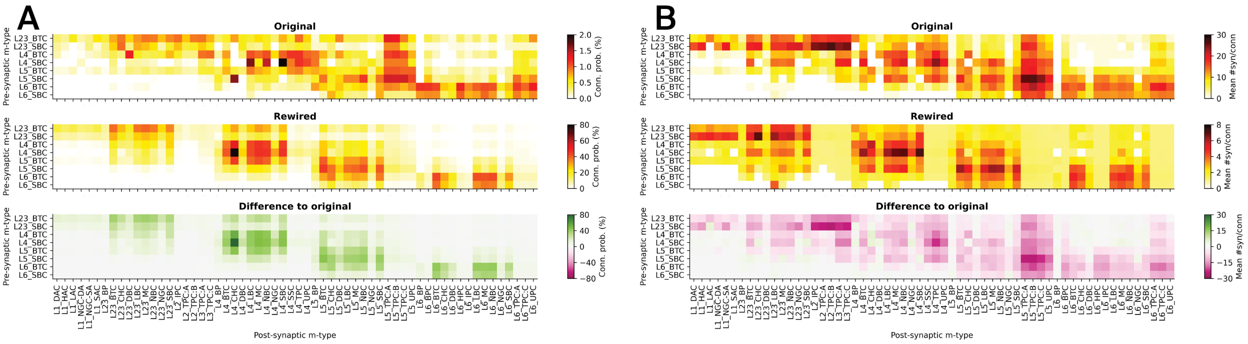


**Figure S1: Detailed changes in rewired VIP+ interneuron connectivity by m-type.**
(A) Mean connection probabilities in the original and rewired connectomes, and the differences between the rewired and original one. Note the different color scales. (B) Same as A, but for the mean number of synapses per connection. These results were obtained by running a structural comparison using the `connectivity` code module (see Table S6).

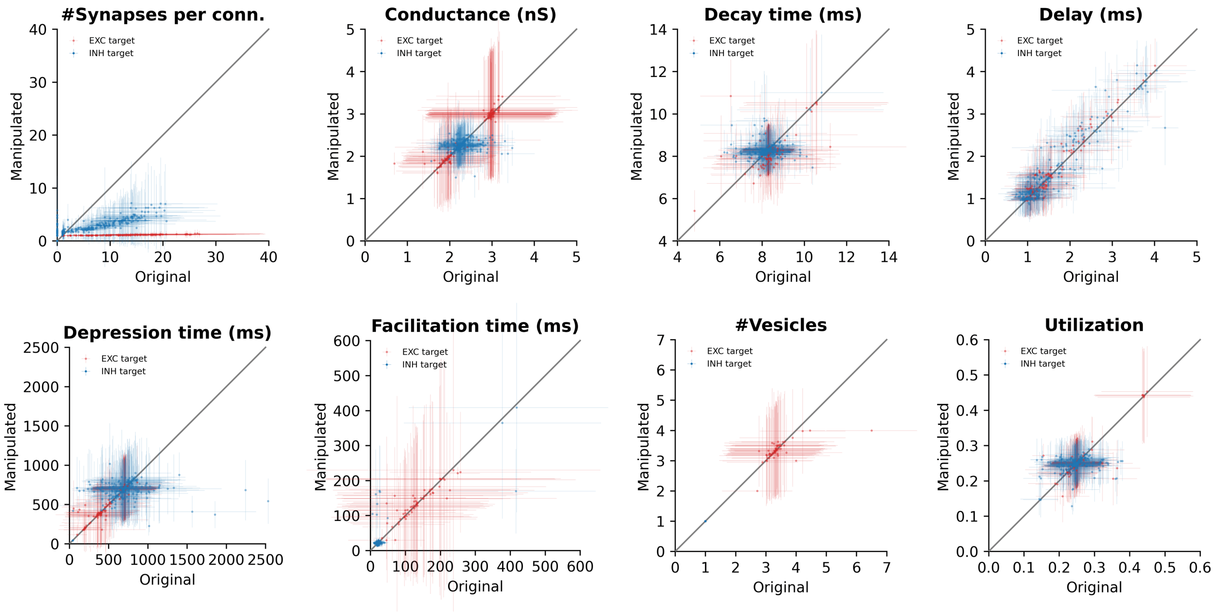


**Figure S2: Validation of synapse parameters in rewired VIP+ interneuron connectivity by m-type.**
Numbers of synapses per connection and important physiological synapse parameter values in the original connectome (x axis) plotted against the manipulated connectome (y axis). Each data point is the mean ± standard deviation of all connections belonging to a pathway, i.e., a pair of $8 \times 60$ pre- and post-synaptic m-types. Post-synaptic m-types are divided into excitatory (18; red) and inhibitory (42; blue) types, as indicated by the legend. These results were obtained by running a structural comparison using the `connectivity` (#Synapses per connection) and `properties` (remaining panels) code modules respectively (see Table S6).

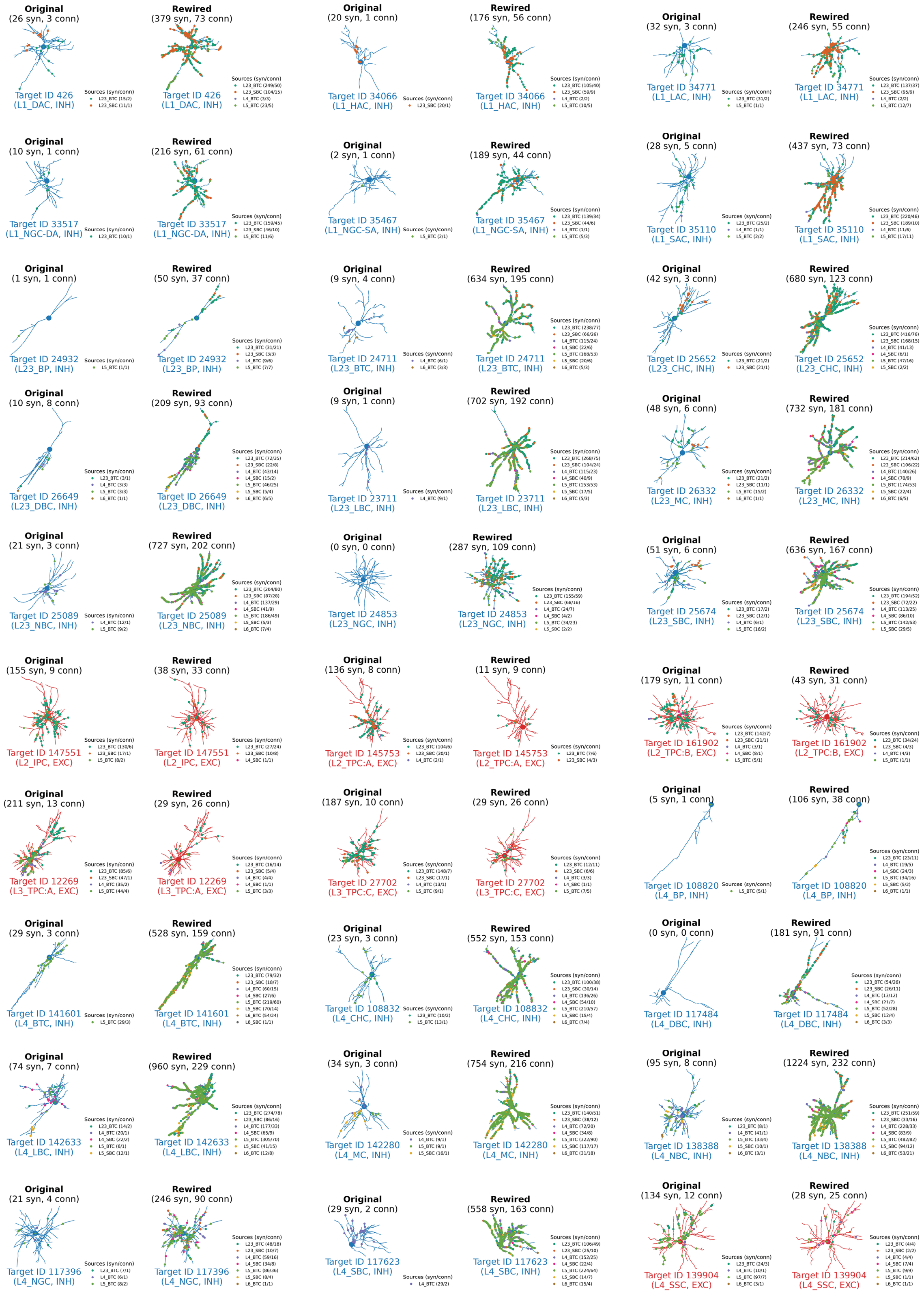


**Figure S3: Examples of synapses on dentritic morphologies before and after VIP+ interneuron rewiring for 30 m-types.**

The dots indicate synapses from different BTC/SBC source types (as indicated by the legend) targeting inhibitory (blue) and excitatory (red) neurons of different m-types in the original vs. the rewired connectome. Small numbers denote numbers of synapses and connections respectively. Example neurons with highest in-degrees were selected. Same style as the examples in Figure 2D. Continued in Figure S4.

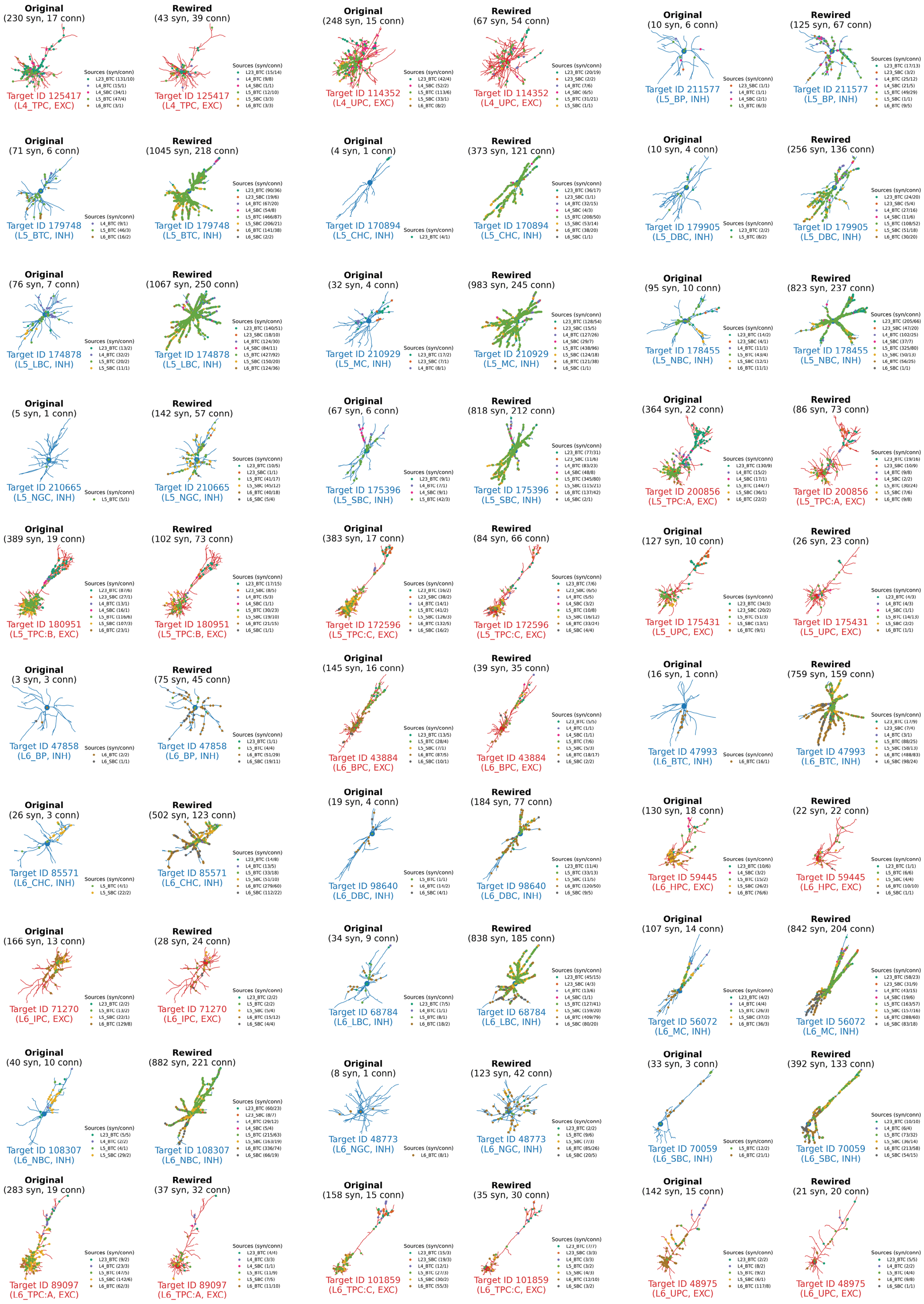


**Figure S4: Examples of synapses on dentritic morphologies before and after VIP+ interneuron rewiring for another 30 m-types.**

The dots indicate synapses from different BTC/SBC source types (as indicated by the legend) targeting inhibitory (blue) and excitatory (red) neurons of different m-types in the original vs. the rewired connectome. Small numbers denote numbers of synapses and connections respectively. Example neurons with highest in-degrees were selected. Same style as the examples in Figure 2D. Continuation from Figure S3.

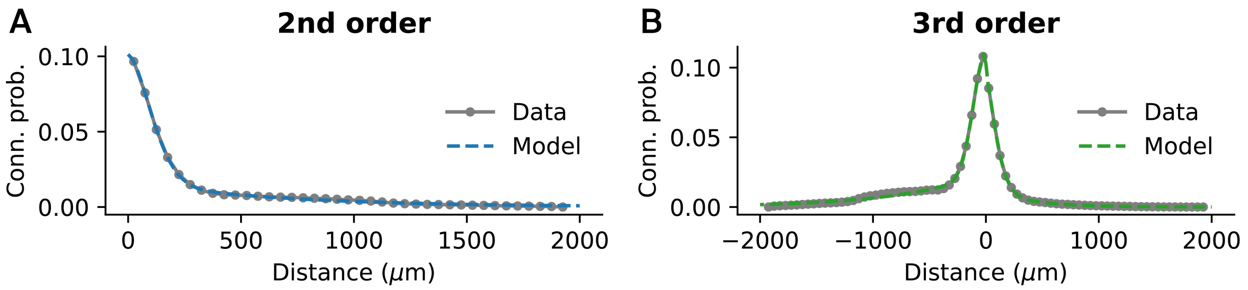


**Figure S5: Extracted data versus stochastic model fits used for simplified connectomes.** Binned distance-dependent connection probabilities extracted from the baseline connectome and connection probabilities obtained from fitted parametric stochastic 2$^{\text{nd}}$ order (A) an 3$^{\text{rd}}$ order (B) connectivity models of types `ConnProb2ndOrderComplexExpModel` and `ConnProb3rdOrderComplexExpModel` respectively. Negative distances in (B) indicate $\Delta z < 0$ whereas positive distances $\Delta z > 0$ (see Eq. S5 in Table S1). More details can be found in the Methods.

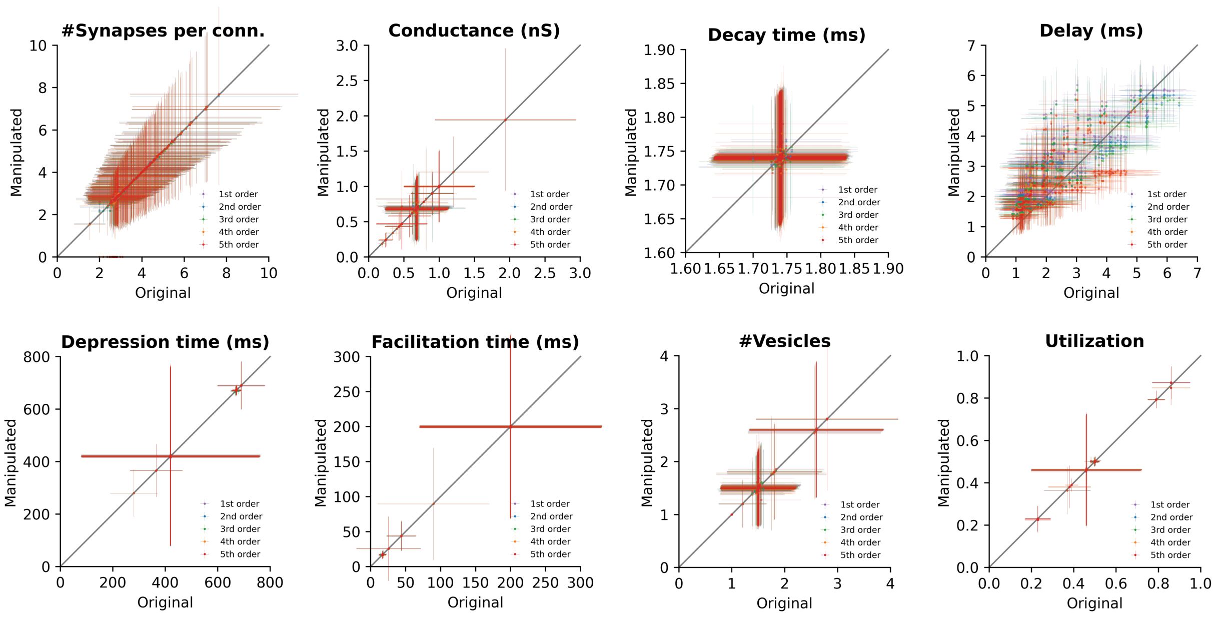


**Figure S6: Validation of synapse parameters in the simplified connectomes by m-type.** Numbers of synapses per connection and important physiological synapse parameter values in the original connectome (x axis) plotted against the simplified connectomes (y axis), as indicated by the legend. Each data point is the mean ± standard deviation of all connections belonging to a pathway, i.e., a pair of 18 × 18 excitatory pre- and post-synaptic m-types (L2_IPC, L2_TPC:A, L2_TPC:B, L3_TPC:A, L3_TPC:C, L4_SSC, L4_TPC, L4_UPC, L5_TPC:A, L5_TPC:B, L5_TPC:C, L5_UPC, L6_BPC, L6_HPC, L6_IPC, L6_TPC:A, L6_TPC:C, L6_UPC). These results were obtained by running structural comparisons using the `connectivity` (#Synapses per connection) and `properties` (remaining panels) code modules respectively (see Table S6).

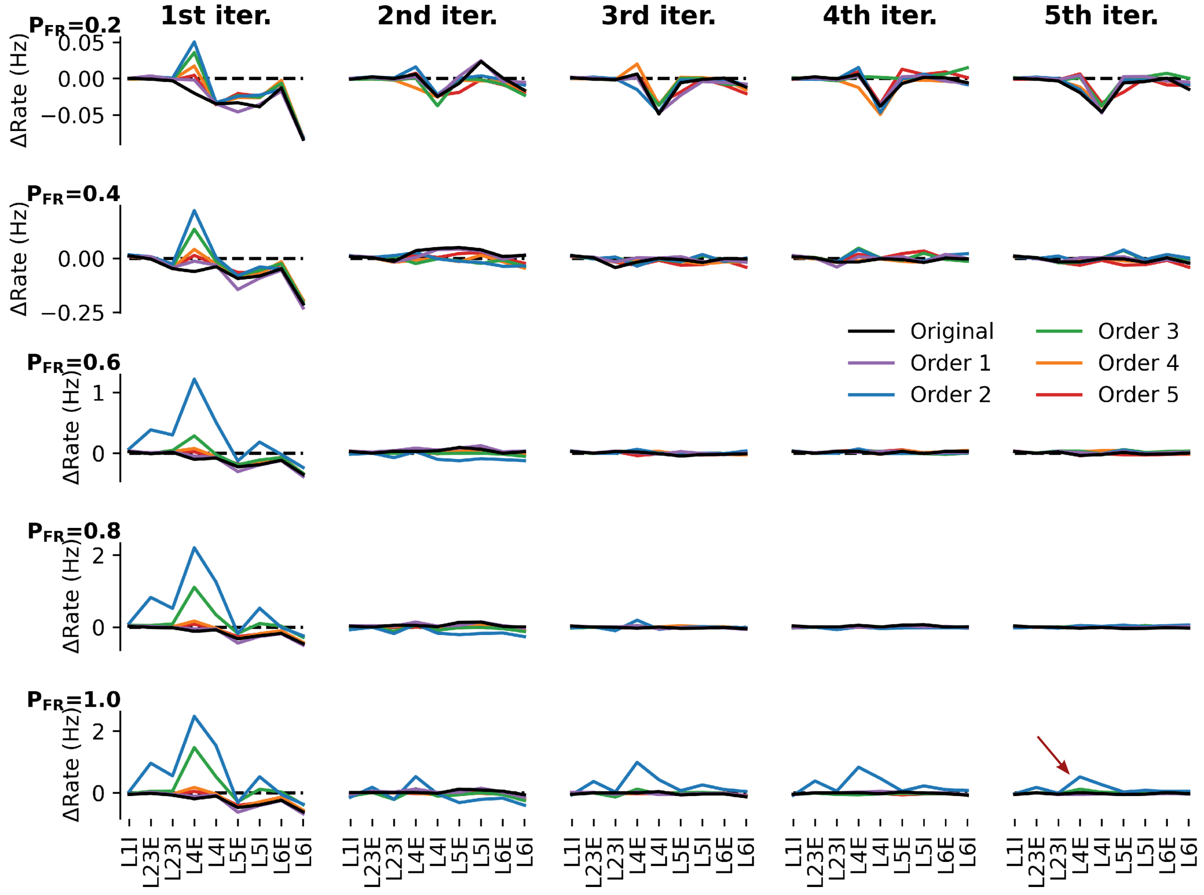


**Figure S7: Spontaneous activity calibration of the simplified connectomes.**
We calibrated population-specific OU-parameters in order to obtain *in vivo*-like spontaneous activity (see Methods) for different fractions $P_{FR}$ (rows) over five iterations (columns). Each individual plot show the firing rate differences ΔRate (y axis) between the observed rates and the fraction $P_{FR}$ of the reference rates for different E/I populations (x axis). The individual colors denote different simplified connectomes as indicated by the legend. Note the different y axis scales. At $P_{FR} = 1.0$, there was still a rate mismatch for the 2$^{nd}$ order connectome even after five iterations (red arrow).

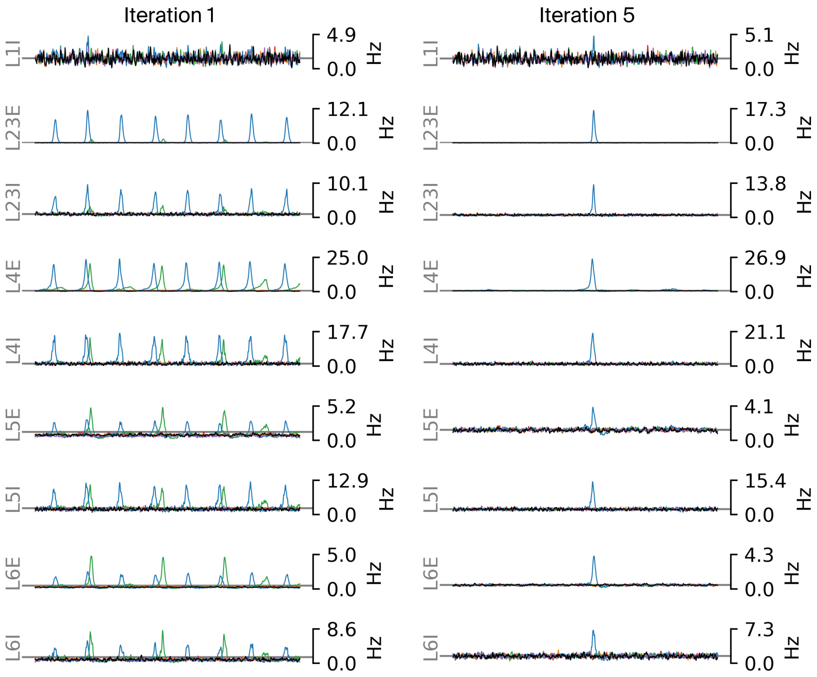


**Figure S8: Network activity before and after recalibration of the simplified connectomes for $P_{FR} = 1.0$.**
Instantaneous firing rates of 5 s spontaneous activity of E and I populations across layers for each of the rewired connectomes before and after recalibration as in Figure 4, but for an *in vivo* fraction $P_{FR} = 1.0$.

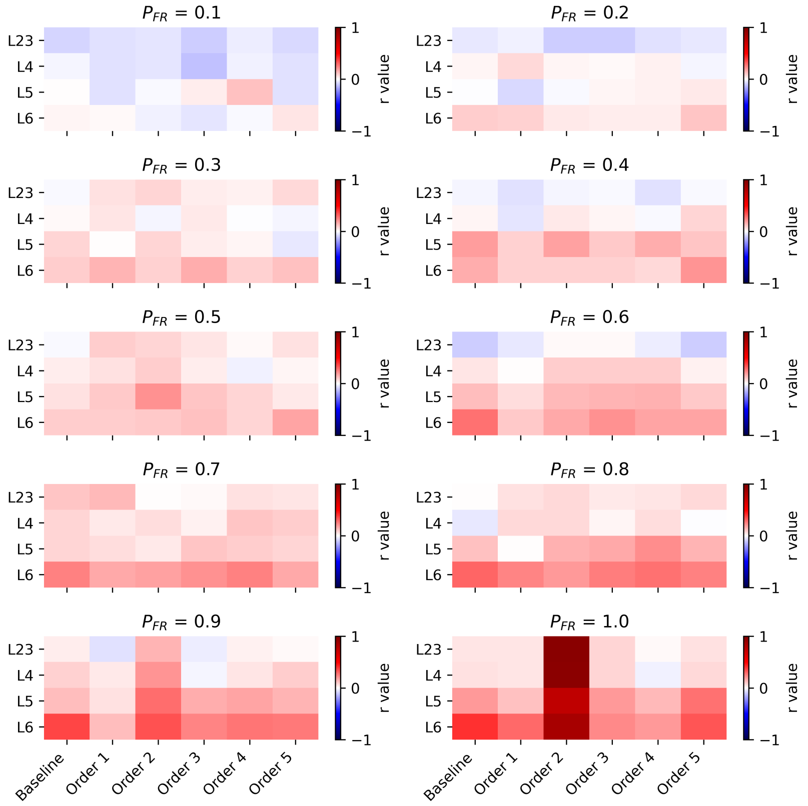


**Figure S9: E/I correlation in the simplified connectomes after recalibration.**
Pearson correlation coefficient (r value) between the instantaneous firing rates (5 ms bin size, Gaussian smoothing with standard deviation 1.0) of 5 s spontaneous activity of the respective E and I populations across layers and different *in vivo* fractions $P_{FR}$.

#### Supplementary Tables

**Table S1: Available stochastic and generic model descriptions and extensions in the connectome manipulation framework.**
The table describes how the different model types are defined in terms of their inputs and outputs. Unless noted otherwise, the pre-synaptic neuron is always indexed with $i$ and the post-synaptic neuron with $j$. The models are grouped by category. Where applicable, the number of associated model parameters is indicated that need to be fitted against data during model building or defined manually when creating a model instance.

**Model & Description**

*Connectivity:*

`ConnProb1stOrderModel`

> $1^{st}$ order (Erdős-Rényi) connection probability model (Gal et al., 2020); returns a constant connection probability
>
> $$P_{ij} = p_{const} \tag{S1}$$
>
> between neurons $i$ and $j$.
> Number of model parameters: 1

`ConnProb2ndOrderExpModel`

> $2^{nd}$ order (distance-dependent) connection probability model (Gal et al., 2020), based on an exponential function; returns the connection probability
>
> $$P_{ij} = f(d_{ij}) = \alpha \cdot e^{-\beta \cdot d_{ij}} \tag{S2}$$
>
> which is a function of the Euclidean (intersomatic) distance $d_{ij}$ between neurons $i$ and $j$.
> Number of model parameters: 2

`ConnProb2ndOrderComplexExpModel`

> $2^{nd}$ order (distance-dependent) connection probability model (Gal et al., 2020), based on a proximal (fast-decaying) and a distal (slowly-decaying) exponential function; returns the connection probability
>
> $$P_{ij} = f(d_{ij}) = \alpha_p \cdot e^{-\beta_p \cdot {d_{ij}}^{\gamma}} + \alpha_d \cdot e^{-\beta_d \cdot d_{ij}} \tag{S3}$$
>
> which is a function of the Euclidean (intersomatic) distance $d_{ij}$ between neurons $i$ and $j$.
> Number of model parameters: 5

`ConnProb3rdOrderExpModel`

> $3^{rd}$ order (bipolar distance-dependent) connection probability model (Gal et al., 2020), based on an exponential function; returns the connection probability
>
> $$P_{ij} = f(d_{ij}, sign(\Delta z)) = \begin{cases} \alpha_- \cdot e^{-\beta_- \cdot d_{ij}}, \text{if } sign(\Delta z) < 0 \\ \alpha_+ \cdot e^{-\beta_+ \cdot d_{ij}}, \text{if } sign(\Delta z) > 0 \\ \text{Mean of both, if } sign(\Delta z) = 0 \end{cases} \tag{S4}$$
>
> which is a function of the Euclidean (intersomatic) distance $d_{ij}$ between neurons $i$ and $j$ and the sign of the difference of their vertical positions (i.e., along z axis; $\Delta z = z_j - z_i$).
> Number of model parameters: 4

`ConnProb3rdOrderComplexExpModel`

$3^{rd}$ order (bipolar distance-dependent) connection probability model (Gal et al., 2020), based on a proximal (fast-decaying) and a distal (slowly-decaying) exponential function; returns the connection probability

$$P_{ij} = f(d_{ij}, sign(\Delta z)) = \begin{cases} \alpha_{p-} \cdot e^{-\beta_{p-} \cdot {d_{ij}}^{\gamma_-}} + \alpha_{d-} \cdot e^{-\beta_{d-} \cdot d_{ij}}, \text{if } sign(\Delta z) < 0 \\ \alpha_{p+} \cdot e^{-\beta_{p+} \cdot {d_{ij}}^{\gamma_+}} + \alpha_{d+} \cdot e^{-\beta_{d+} \cdot d_{ij}}, \text{if } sign(\Delta z) > 0 \\ \text{Mean of both, if } sign(\Delta z) = 0 \end{cases} \quad \text{(S5)}$$

which is a function of the Euclidean (intersomatic) distance $d_{ij}$ between neurons $i$ and $j$ and the sign of the difference of their vertical positions (i.e., along z axis; $\Delta z = z_j - z_i$).
Number of model parameters: 10

`ConnProb4thOrderLinInterpnModel`

$4^{th}$ order (offset-dependent) connection probability model (Gal et al., 2020), based on multidimensional linear interpolation of probability values given on a regular grid of x, y, and z offsets; returns the connection probability

$$P_{ij} = f(\Delta x, \Delta y, \Delta z) \quad \text{(S6)}$$

which is a function of the x, y, and z offsets between neurons $i$ and $j$ (i.e., $\Delta x = x_j - x_i$, $\Delta y = y_j - y_i$, $\Delta z = z_j - z_i$).
Number of model parameters: #Grid points (i.e., bins)

`ConnProb4thOrderLinInterpnReducedModel`

$4^{th}$ order (offset-dependent) connection probability model (Gal et al., 2020), based on multidimensional linear interpolation of probability values given on a regular grid of radial (r; within x/y plane) and axial (z) offsets; returns the connection probability

$$P_{ij} = f(\Delta r, \Delta z) \quad \text{(S7)}$$

which is a function of the offsets in radial and axial direction between neurons $i$ and $j$ (i.e., $\Delta r = \sqrt{\Delta x^2 + \Delta y^2}$, $\Delta z = z_j - z_i$).
Number of model parameters: #Grid points (i.e., bins)

`ConnProb5thOrderLinInterpnModel`

$5^{th}$ order (position-dependent) connection probability model (Gal et al., 2020), based on multidimensional linear interpolation of probability values given on a regular grid of x, y, and z positions and offsets; returns the connection probability

$$P_{ij} = f(x_i, y_i, z_i, \Delta x, \Delta y, \Delta z) \quad \text{(S8)}$$

which is a function of the x, y, and z positions of the pre-synaptic neuron $i$ and offsets between neurons $i$ and $j$ (i.e., $\Delta x = x_j - x_i$, $\Delta y = y_j - y_i$, $\Delta z = z_j - z_i$).
Number of model parameters: #Grid points (i.e., bins)

`ConnProb5thOrderLinInterpnReducedModel`

$5^{\text{th}}$ order (position-dependent) connection probability model (Gal et al., 2020), based on multidimensional linear interpolation of probability values given on a regular grid of axial (z) positions and radial (r; within x/y plane) and axial offsets; returns the connection probability

$$P_{ij} = f(z_i, \Delta r, \Delta z) \tag{S9}$$

which is a function of the axial position of the pre-synaptic neuron $i$ and the offsets in radial and axial direction between neurons $i$ and $j$ (i.e., $\Delta r = \sqrt{\Delta x^2 + \Delta y^2}$, $\Delta z = z_j - z_i$).
Number of model parameters: #Grid points (i.e., bins)

`ConnProbAdjModel`

Connection probability model, defined by a sparse adjacency matrix; returns the connection probability 1 if a connection between neurons $i$ and $j$ exists, otherwise 0, i.e., this is essentially a deterministic connectivity representation.
Number of model parameters: #Connections

*Physiology:*

`LinDelayModel`

Linear distance-dependent axonal delay model, optionally with pathway-specific model attributes for different combinations of pre-/post-synaptic m-types; returns randomly drawn delay values from a truncated normal distribution with given mean

$$\mu_{ij} = \alpha_D + \beta_D * d_{ij}, \tag{S10}$$

standard deviation $\sigma_D$ (constant), and minimum delay $D_{min}$ (constant), i.e., the mean delay $\mu_{ij}$ linearly depends on the Euclidean distance $d_{ij}$ between the soma of a pre-synaptic neuron $i$ and the synapse position on the post-synaptic dendrite of neuron $j$.
Number of model parameters: 2

`ConnPropsModel`

Connection properties model for realizing connections by forming synapses, based on individual parameters for pathways, i.e., pairs of pre-/post-synaptic m-types; returns synapse properties (e.g., conductance, time constants, readily-releasable vesicles, etc.) drawn from given distributions, such as constant, (truncated) normal, gamma, (zero-truncated) Poisson, or discrete distributions, for a single connection. Each distribution is defined by a specific set of attributes, e.g., mean and standard deviation for a normal distribution. A full list of distribution attributes can be found in the Documentation (under "Model fitting API"). The numbers of synapses per connection are either drawn from a given distribution or provided externally. The same property values can be shared among all synapses belonging to the same connection; otherwise, they are drawn independently. This model type is not restricted to a predefined set of synapse properties; it may contain distributions for any number of properties that are specified by the user.
Moreover, correlations between synaptic property values can optionally be specified by means of a covariance matrix. For generating correlated parameter values, we implemented the method outlined in Chindemi et al. (2022). If specified, parameter values are jointly drawn from a multivariate normal distribution with zero mean and given covariance matrix (with diagonal entries one). These values are then remapped to the respective marginal distributions as defined in the model. A minimal working example can be found in the `/examples` folder in the GitHub repository.
Number of model parameters: #Pathways × #Synapse properties × #Distribution attributes + #Pathways × #Correlated pairs of properties (optional)

*Model extensions:*

`PosMapModel`

> Position mapping from one coordinate system to another for a given set of neurons; returns the mapped soma position for a given neuron ID. This model extension can be used in combination with all connection probability models depending on spatial geometry, e.g., for aligning the coordinate axes with the cortical layers using a flat/depth map projection.
> Importantly, when using a position mapping for fitting a connection probability model, the same position mapping must be used used when applying the connection probability model, e.g., for (re)wiring.

*Generic:*

`LookupTableModel`

> Generic lookup table to access any (sparse) information based on pre- and post-synaptic neuron IDs; returns zero if no value is stored for a given pair of neurons. For example, it can be used to specify adjacency matrices (boolean entries), connection probabilities between neurons (decimal numbers between 0.0 and 1.0), or synaptome matrices (integer numbers of synapses per connection).

`PropsTableModel`

> Generic properties table to store a table of properties based on pre- and post-synaptic neuron IDs; it may contain multiple entries for a given pair of neurons. For example, it can be used to store exact synapse positions precomputed externally.

**Table S2: Model building tools in the connectome manipulation framework.**
The table describes the available tools (Python modules) for extracting data, fitting stochastic models, and storing them in a model representation format. All tools work in three steps: data extraction, model fitting (where applicable) and storing, and visualization of the actual data and/or model output. Some of the tools optionally support the use of cross-validation in order to prevent overfitting (see Methods).

**Model & Description**

---

*Connectivity:*

`conn_prob.py`

> Model fitting of stochastic connection probability models with given model order (`ConnProb...Order` models in Table S1), based on binned connection probabilities (with given bin sizes) extracted from an existing connectome. Cross-validation is supported when fitting such models. Also, bins with too few data points, which would usually result in noisy probability estimates, can be excluded by setting a lower threshold. For the (parametric) $2^{nd}$ and $3^{rd}$ order models, we use least squares for parameter fitting (`scipy.optimize.curve_fit` function); an optional relative error threshold can be specified to prevent building a model from a bad fit. For the (non-parametric) $4^{th}$ and $5^{th}$ order models, the binned probability estimates can optionally be smoothed by a multidimensional Gaussian filter (`scipy.ndimage.gaussian_filter` function) to reduce noise before building the model.

`conn_prob_adj.py`

> Adjacency matrix extraction from an existing connectome and storing it as `ConnProbAdjModel` (Table S1).

*Physiology:*

`delay.py`

Axon delay model fitting of type `LinDelayModel` (Table S1) based on binned distance-dependent synaptic delays (with given bin size) between samples of neurons extracted from an existing connectome. Least squares linear regression (`sklearn.linear_model.LinearRegression` function) is used to determine the model coefficient $\alpha_d$ and $\beta_d$ for the mean delay (Eq. S10); its standard deviation $\sigma_D$ is determined as the mean of the standard deviations of all bins, $D_{min}$ as the minimum overall delay. Cross-validation is supported when fitting such a model.

`conn_props.py`

Connection and synapse properties model fitting of type `ConnPropsModel` (Table S1) for pairs of pre-/post-synaptic m-types. Statistics for connections (in case of shared values among synapses) or individual synapses between samples of neurons are extracted from an existing connectome for each pair of m-types and used to parameterize the individual model distribution types specified for each property. Distribution fitting is not restricted to a specific set of properties; any properties that are present in a given connectome may be included. Cross-validation is supported when fitting such a model.

Missing values for pairs of m-types (i.e., between which there are no or too few connections to estimate reliable statistics) are gradually interpolated at different levels of granularity from similar pathways with matching pre-/post-synaptic m-type, layer, or E/I cell type, if available:

| Level | Pre-synaptic | Post-synaptic |
|---|---|---|
| 0 | m-type | layer & cell type |
| 1 | m-type | cell type |
| 2 | layer & cell type | layer & cell type |
| 3 | cell type | cell type |
| 4 | any | any |

*Model extensions:*

`pos_mapping.py`

Building a position mapping extension of type `PosMapModel` (Table S1) from a voxelized brain atlas in NRRD (Nearly Raw Raster Data) format consisting of a flat map (x/y plane, parallel to cortical layers) and a cortical depth map (z axis, perpendicular to cortical layer). The soma positions of a given population are transformed to the new coordinate system by linear interpolation between voxel positions, if possible; otherwise, nearest-neighbor interpolation is employed.

`pos_mapping_from_table.py`

Building a position mapping extension of type `PosMapModel` (Table S1), by just loading the transformed soma positions for a given population from a position table precomputed externally.

**Table S3: Manipulation operations in the connectome manipulation framework.**
The table describes the available manipulation operations (Python modules) and how they can be configured. The modules are grouped by types of use cases as in Figure 1C.

**Module & Description**

*Testing:*

`null_manipulation.py`

Dummy manipulation not performing any manipulation at all, i.e., the output is identical to the input connectome. This operation is intended as a test condition to run the manipulation pipeline without actually manipulating the connectome.

*Remove synapses:*

`syn_removal.py`

Removes a certain percentage of randomly sampled synapses between selected populations of neurons. Optionally, connections can be kept (i.e., keeping at least 1 synapse per connection), and rescaling of synaptic conductances is possible in order to keep the sum of conductances per connection constant.

`syn_subsampling.py`

Random subsampling of synapses, keeping a certain percentage of randomly selected synapses. This is a special case of the functionality implemented in `syn_removal.py`, which is more efficient but does not have any selection options supported.

*Alter synapses:*

`syn_prop_alteration.py`

Alters values of selected synapse properties (e.g., conductance, delay, ...) of a certain percentage of randomly sampled synapses between selected populations of neurons. Values can be altered according to the following options:

| Option | New values given by |
|---|---|
| `setval` | Absolute value |
| `scale` | Relative scaling by a given value |
| `offset` | Offsetting by a give value |
| `shuffle` | Shuffling among synapses |
| `randval` | Random absolute value drawn from a given distribution |
| `randscale` | Relative scaling by a value drawn from a given distribution |
| `randadd` | Adding a random value drawn from a given distribution |

This operation can be utilized in various applications for investigating the impact of certain synapse properties (e.g., lower/higher conductances, shorter/longer delays, etc.) on the emerging network activity. It can also be used to (carefully) manipulate synapse locations on the dendrite, e.g., placing them on the soma, by setting their section index and offset to 0.

*Remove connections:*

`conn_removal.py`

Removes a certain percentage of randomly sampled connections (i.e., all synapses belonging to a connection) between selected populations of neurons. Optionally, only connections within a certain range of synapses per connection, and/or within a specific connection mask can be removed.

A potential application of this manipulation is to systematically study the impact of missing connections on the network activity, as in lesion experiments.

`conn_extraction.py`

Extracts the connectome of a given node set (i.e., a named sub-population of neurons), keeping only connections between neurons within that node set and removing all connections from, to, and between neurons outside that node set. Node sets intrinsic to a circuit or provided through an external SONATA node sets file (JSON format) containing a list of neuron IDs are supported.

Potential applications are restricting the connectivity to certain regions of interest, or as pre-processing step to reduce the connectome size in order to obtain a new baseline connectome for further manipulations.

*Rewire, transplant, wire:*

`conn_rewiring.py`

Fundamental operation for (re)wiring connections between populations of neurons, which involves removing and changing existing, as well as creating new connections. Connectivity is defined by a given connection probability model (`ConnProb...` models in Table S1) with or without use of a position mapping (`PosMapModel`), which can optionally be scaled by a global probability scaling factor `p_scale` to fine-tune the resulting number of connections. Existing connections can be preserved or reused in new connections; existing synapses (and connections) may be deleted and new ones created by either reusing existing or randomly generating new positions on the post-synaptic dendrites, or loading existing positions externally. Also, in-degrees can be preserved, and rewiring can be restricted to only adding or deleting connections; see Table S4 for all options. Importantly, (re)wiring operations are restricted to outgoing connections from either excitatory or inhibitory neuron types at a time, to prevent intermixing of different synapse classes when reusing synapses in new connections, which would be a violation of Dale's law (Strata & Harvey, 1999).

Physiological synapse properties can be sampled from existing synapses or drawn from pathway-specific model distributions (`ConnPropsModel` in Table S1); synaptic delays can be assigned depending on the distance to their pre-synaptic neurons (`LinDelayModel`). Numbers of synapses per connection can be sampled from existing connections, drawn from model distributions, or provided by a synaptome matrix (stored as `LookupTableModel`).

The main applications of this operation are to create connectivity for network models without connectome, and to investigate the impact of rewired or transplanted connectivity in existing connectomes.

**Table S4: Rewiring options.**
The table describes the available options for (re)wiring (i.e., by use of `conn_rewiring`) and how they affect the resulting connectivity. Depending on these options, certain aspects of the existing connectivity can be preserved, as summarized in Table S5. In case of wiring an empty connectome from scratch, only rewiring options that do not involve an existing connectome are applicable, as indicated by the footnote.

| **Option & Description** |
|---|
| *Connections:* |
| `keep_conns` |
| Keeping existing connections. If during the connectivity assignment step (see Figure 1D1) connections are to be established that already exist in the input connectome, such connections are kept exactly as they are. That is, the pre-synaptic neuron (i.e., connection source), number of synapses per connection, synapse positions on the post-synaptic dendrite, as well as synapse physiology are preserved. |
| `reuse_conns` |
| Reusing existing connections. During rewiring, existing connections are reused to realize new connections in the rewired connectome. Specifically, synapses per connection, synapse positions on the post-synaptic dendrite as well as synapse physiology are preserved, but new pre-synaptic neurons are randomly assigned to these connections. If the number of incoming connections to be established with a post-synaptic neuron exceeds the number of existing (incoming) connections, all existing ones are reused and the remaining ones created as in the general rewiring case. If the number of existing connections is higher, a subset of them are randomly selected and reused, and the remaining ones are deleted. |
| `keep_conns & reuse_conns` |
| In case both options are selected, the behavior will be like `keep_conns` when establishing connections that already exist in the input connectome. For connections that exist in the input but no longer in the rewired connectome, the behavior is like `reuse_conns`, i.e., such connections may be reused to establish other new (incoming) connections. |
| Otherwise† |
| Disregarding existing connections. In the general rewiring case, existing connections are neither kept nor reused but replaced by newly generated (incoming) connections. |
| *Synapse physiology:* |
| `sample` |
| Generation method for physiological parameterization based on sampling. When creating new synapses to form new connections, the number of synapses per connection as well as their physiological property values are randomly sampled from other existing synapses. The existing synapses to sample from are always of the same synapse class as the new synapses that are to be created (i.e., either excitatory or inhibitory). Also, only synapses on post-synaptic neurons belonging to the same m-type as the post-synaptic target neuron are considered. If such synapses don't exist, synapses of the same layer (but non-matching m-type) are considered. If such synapses don't exist either, synaptic properties are sampled from synapses of all available post-synaptic neurons (i.e., neither matching m-type nor layer). When rewiring is run with more than one data split (see Figure 5), property values can only be sampled from synapses within the same split, in which case the `randomization` method is highly recommended which is independent of data splits. |

`randomize`†

Generation method for physiological parameterization based on randomization. When creating new synapses to form new connections, their physiological property values are randomly drawn from given (pathway-specific) property distributions (Table S1). Numbers of synapses per connection are either randomly drawn as well or can be provided through a synaptome matrix.

*Synapse positions:*

`reuse`

Reusing existing synapse positions. Existing synapse positions on the post-synaptic dendrite are reused when generating new synapses. Specifically, for each new connections to be created, the corresponding number of synapses are randomly sampled from all existing synapses, and new synapses are placed at their exact positions. For multi-synaptic connections, the synapse positions are drawn without replacement, if possible. The advantage of reusing positions is that their overall distribution of positions is preserved (but not for a single connection), and that rewiring runs faster since no access to dendritic morphologies is required. Potential drawbacks are that pathway-specific dendritic targeting preferences are not respected, and that multiple synapses belonging to the same or different connections may be placed at exactly the same position, especially for post-synaptic neurons with few existing synapses.

`reuse_strict`

Restricted reuse of existing synapse positions. Same as `reuse`, but reusing only existing synapses that are incoming from the selected source population of neurons. In this way, a pathway-specific dendritic targeting preference can be preserved.

`randomize`†

Randomizing synapse positions. New positions are randomly drawn based on the actual dendritic morphologies. Specifically, each new position is uniformly drawn from the soma and all dendritic morphology sections, and a random (relative) offset within each section (except soma, which has zero offset by definition). This method has lower risk of duplicate synapse positions, but is slower since access to dendritic morphologies and recomputation of 3D synapse positions based on their drawn sections and offsets are required.

`external`†

Loading external synapse positions. New positions are directly loaded from an external position table (provided as `PropsTableModel`, see Table S1). An error is raised if not enough positions are available for a given connection. In case the table contains more positions than required, they are sequentially loaded. This option should be used with caution, since external positions are loaded without consistency checks against actual morphologies.

*In-degree:*

`keep_indegree`

Keeping the in-degrees constant. The number of incoming connections for each post-synaptic neuron which is subject to rewiring is preserved. This is achieved in the connectivity assignment step (see Figure 1D1), by drawing exactly the same number of pre-synaptic neurons as in the input connectome to be connected with a post-synaptic neuron.

This imposes certain constraints on the rewiring operation: (i) The connection probabilities obtained from a stochastic connection probability model (see Table S1) are not interpreted in absolute but in relative terms, so that the overall distribution of connection probabilities may not be fulfilled exactly, but only for each post-synaptic neuron independently. (ii) At least the required number of pre-synaptic neurons must have non-zero connection probabilities to be connected with a post-synaptic neuron, otherwise keeping the in-degree is not possible. (iii) Keeping the in-degree is not compatible with rewiring options that allow only adding or only deleting connections.

Otherwise†

Disregarding existing in-degrees. The numbers of incoming connections for each post-synaptic neuron are not guaranteed to be preserved. Instead, any number of pre-synaptic neurons may be assigned to be connected with a post-synaptic neuron based on the connection probabilities obtained from a stochastic connection probability model (see Table S1).

*Restricted rewiring:*

`add_only`†

During connectivity assignment (see Figure 1D1), connections can be only added; no existing connections will be deleted (but may be reparameterized depending on the other options selected). Hence, the resulting number of incoming connections to a post-synaptic neuron is always greater than or equal to the respective number in the input connectome.

`delete_only`

During connectivity assignment (see Figure 1D1), connections can be only deleted; no new connections will be established. Hence, the resulting number of incoming connections to a post-synaptic neuron is always less than or equal to the respective number in the input connectome.

Otherwise†

No restrictions on rewiring; that is, new connections may be established and existing ones deleted.

*Matching total number of connections (see Methods):*

`p_scale`

Global scaling factor for adjusting the connection probabilities given by a connectivity model (see Table S1).

`estimation_run`

Rewiring operation with early stopping, which does not generate an actual connectome or output file, but writes an estimate of the average number of incoming connections for each post-synaptic neuron that is subject to rewiring into a data log file.

`opt_nconn`

The drawn number of incoming connections during connectivity assignment (see Figure 1D1) in a single random instance of the connectome will be optimized to match its expected number of connections on average.

† Applicable for wiring an empty connectome from scratch.

**Table S5: Preserved aspects of connectivity in rewired connectomes.** The table shows different groups of options (Table S4) regarding existing connections (blue shaded), synapse physiology (green shaded), synapse positions (red shaded), and in-degree (yellow shaded) that can be used for rewiring (i.e., by use of `conn_rewiring`) and how they affect certain aspects of connectivity in the rewired connectome. The table entries indicate whether or not a given property is preserved for a post-synaptic neuron (Y: yes, N: no), or if it is independent of a certain option (n/a: not applicable).

| | | Preserved properties | | | | |
|---|---|---|---|---|---|---|
| **Option** | **Description** | Conn. source | Syn. per conn. | Syn. physiol. | Syn. position | In-degree |
| *Connections* | | | | | | |
| `keep_conns`* | Keeping existing connections | Y | Y | Y | Y | n/a |
| `reuse_conns`* | Reusing existing connections | N | Y | Y | Y | n/a |
| Otherwise | | N | n/a | n/a | n/a | n/a |
| *Synapse physiology* | | | | | | |
| `sample` | Sampling from existing synapses | n/a | N | N | n/a | n/a |
| `randomize` | Randomly drawing from distributions | n/a | N | N | n/a | n/a |
| *Synapse positions* | | | | | | |
| `reuse<_strict>` | Reusing existing synapse positions | n/a | n/a | n/a | Y | n/a |
| `random` | Randomly generating synapse positions | n/a | n/a | n/a | N | n/a |
| `external` | Externally loading synapse positions | n/a | n/a | n/a | N | n/a |
| *In-degree* | | | | | | |
| `keep_indegree` | Keeping the in-degrees constant | n/a | n/a | n/a | n/a | Y |
| Otherwise | | n/a | n/a | n/a | n/a | N |

* If both are selected, `keep_conns` will override `reuse_conns`.

**Table S6: Tools for structural comparison in the connectome manipulation framework.** The table describes the available tools (Python modules) for comparing connectomes in terms of connectivity structure and synaptic physiology. All tools work in two steps: computing metrics to compare, and visualizing them for the two connectomes and their difference.

| **Module & Description** |
|---|
| *Connectivity:* |
| `adjacency.py` |
| Structural comparison of two connectomes in terms of adjacency and synaptome matrices (i.e., connectivity and synapse counts between pairs of individual pre-/post-synaptic neurons) for selected pathways. |
| `connectivity.py` |
| Structural comparison of two connectomes in terms of connection probability and mean number of synapses per connection between groups of neurons (grouped by a given cell property, e.g., by layer, m-types, etc.) for selected pathways. |
| *Physiology:* |
| `properties.py` |
| Comparison of two connectomes in terms of statistical properties (e.g., mean, standard deviation, etc.) of selected synapse parameters of connections between groups of neurons (grouped by a given cell property, e.g., by layer, m-types, etc.) for selected pathways. |

### References


Chindemi, G., Abdellah, M., Amsalem, O., Benavides-Piccione, R., Delattre, V., Doron, M., ... Muller, E. B. (2022). A calcium-based plasticity model for predicting long-term potentiation and depression in the neocortex. *Nature Communications*, *13*(1), 3038. doi: 10.1038/s41467-022-30214-w

Gal, E., Perin, R., Markram, H., London, M., & Segev, I. (2020). Neuron geometry underlies universal network features in cortical microcircuits. *bioRxiv*. doi: 10.1101/656058

Strata, P., & Harvey, R. (1999). Dale's principle. *Brain Research Bulletin*, *50*(5), 349–350. doi: 10.1016/S0361-9230(99)00100-8
