## Supplementary material for "A connectome manipulation framework for the systematic and reproducible study of structure–function relationships through simulations": List of technical terms

**Adjacency matrix** Matrix representing whether or not there is a connection between any given pair of pre- and post-synaptic neurons.

**Connectome** Comprehensive structural description of the synaptic connections between neurons within the brain or individual brain regions.

***In silico*** Experiments carried out on a computer by means of a simulation software.

**Network function** Particular spiking activity of the neurons in a neural network generated spontaneously or in response to external stimulation.

**Network structure** Organization of synaptic connectivity among neurons in a neural network.

**Pathway** Entirety of connections between specific pre- and post-synaptic populations of neurons selected based on criteria such as their morphological type (m-type).

**Recalibration** Iterative parameterization procedure for layer-specific conductance injections into neurons until their firing rates match expected values.

**Rewiring** Manipulation of synaptic connections between populations of neurons by creating new connections and removing or changing existing ones.

**Stochastic model** Probabilistic mathematical description of certain aspects of connectivity, e.g., connection probabilities or synaptic parameter distributions.

**Synaptome matrix** Matrix representing the number of synapses that form a connection between any given pair of pre- and post-synaptic neurons.

**Transplanting** Transfer of physiological and/or structural connectivity characteristics of single connections, pathways, or entire connectomes from one to another connectome.

**Wiring** Generation process of a connectome from scratch by creating new synaptic connections between (existing) populations of neurons.
